# Apolipoprotein E abundance is elevated in the brains of individuals with Down syndrome-Alzheimer’s disease

**DOI:** 10.1101/2025.02.24.639862

**Authors:** Clíona Farrell, Yazead Buhidma, Paige Mumford, Wendy E. Heywood, Jenny Hällqvist, Lisi Flores-Aguilar, Elizabeth Andrews, Negin Rahimzadah, Orjona Stella Taso, Eric Doran, Vivek Swarup, Elizabeth Head, Tammaryn Lashley, Kevin Mills, Christina E. Toomey, Frances K. Wiseman

**Affiliations:** UK Dementia Research Institute at University College London; London, United Kingdom; Queen Square Institute of Neurology, University College London; London, United Kingdom; UCL Great Ormond Street Institute of Child Heath, University College London, London, United Kingdom; Department of Pathology and Laboratory Medicine, University of California, Irvine, CA, USA; Mathematical, Computational, and Systems Biology (MCSB) Program, University of California, Irvine, Irvine, CA, USA; Institute for Memory Impairments and Neurological Disorders (MIND), University of California, Irvine, Irvine, CA, USA; Center for Complex Biological Systems (CCBS), University of California Irvine, Irvine, CA, USA; Department of Neurobiology and Behaviour, University of California, Irvine, CA, USA; Department of Pediatrics, University of California, Irvine, School of Medicine, Orange, CA, USA; The Francis Crick Institute, London, United Kingdom

**Keywords:** Trisomy 21, ApoE, Amyloid Precursor protein, mass spectrometry, neuropathology, frontal cortex

## Abstract

Trisomy of chromosome 21, the cause of Down syndrome (DS), is the most commonly occurring genetic cause of Alzheimer’s disease (AD). Here, we compare the frontal cortex proteome of people with Down syndrome-Alzheimer’s disease (DSAD) to demographically matched cases of early-onset AD and healthy ageing controls. We find wide dysregulation of the proteome, beyond proteins encoded by chromosome 21, including an increase in the abundance of the key AD-associated protein, APOE, in people with DSAD compared to matched cases of AD. To understand the cell types that may contribute to changes in protein abundance, we undertook a matched single-nuclei RNA-sequencing study, which demonstrated that *APOE* expression was elevated in subtypes of astrocytes, endothelial cells and pericytes in DSAD. We further investigate how trisomy 21 may cause increased APOE. Increased abundance of APOE may impact the development of, or response to, AD pathology in the brain of people with DSAD, altering disease mechanisms with clinical implications. Overall, these data highlight that trisomy 21 alters both the transcriptome and proteome of people with DS in the context of AD, and that these differences should be considered when selecting therapeutic strategies for this vulnerable group of individuals who have high-risk of early-onset dementia.

## Introduction

Down syndrome (DS) is a common genetic condition caused by an extra copy of human chromosome 21 (Hsa21) (trisomy 21), which contains approximately 221 protein-coding genes (Ensembl release 109 – Feb 2023). In the UK, there are approximately 47,000 individuals with DS. By age 40 years, most individuals with DS will have developed Alzheimer’s disease (AD) pathology, characterised by the accumulation of amyloid-β plaques and neurofibrillary tau tangles (NFT) within the brain [17, 67]. By the age of 55 years, more than half of people with DS will have developed clinical dementia, caused by AD, and by age 70 years, virtually all individuals with DS will have AD-dementia [27, 32, 39, 44]. Down syndrome-Alzheimer’s disease (DSAD) is the most commonly occurring single genetic cause of AD, and is a result of triplication of the amyloid precursor protein gene (*APP*) on Hsa21, which results in the increased production of the amyloid-β peptide and other *APP* gene products [11, 51]. DSAD shares many features of AD in the general population, but the additional copy of Hsa21 alters cellular and molecular processes, resulting in differences in some aspects of the disease. These include a faster development of tau pathology in relation to amyloid-β [71], changes to neuroimmune biology [48, 43, 65], and increased incidence of seizures [2]. These differences in AD development occur because trisomy 21 results in an imbalance of Hsa21 gene products, leading to a modification of cell state and function; but how specific Hsa21-encoded genes cause these changes is not currently known. Evidence in preclinical models demonstrates that genes other than *APP* being in three-copies modifies aspects of DSAD, such as cognition and amyloid-β load [68, 1, 47], but more work is required to understand how the molecular processes of AD are altered by trisomy 21 in the brain of people with DS.

Recent transcriptomic studies have begun to address this by systematically interrogating how the extra copy of Hsa21 alters gene expression within the brain of people with DS, demonstrating changes in the inhibitory to excitatory neuronal ratio, and microglial cell states [46, 48]. These studies have not fully addressed how transcriptomic signatures in the brain of people with DSAD compare to AD in the general population, leaving the effects of trisomy 21 and AD on molecular processes still to be disentangled. Moreover, the functional effects of gene dosage imbalance are often mediated at the protein level. Homeostatic processes can regulate the abundance of proteins such that the raised level of a transcript does not always increase the abundance of the corresponding protein [7, 53, 56]. Whether the brain proteome of people with DSAD is also altered alongside the transcriptome has not yet been studied.

We hypothesised that as well as the transcriptome, trisomy 21 alters the proteome of the brain of people with DSAD, and that it differs compared to the AD-associated proteome from the general population. To address this, in this study, we conducted a matched proteomic and single-nuclei transcriptomic study comparing cases of DSAD with age and demographically matched cases of early-onset AD (EOAD) and healthy ageing (HA) euploid individuals.

## Materials and Methods

### Human post-mortem tissue ethics statement

Use of human post-mortem brain tissue in this study was carried out in accordance with the Human Tissue Act (2004). Samples were received from both Newcastle Brain Tissue Resource (NBTR), Newcastle University, UK; South West Dementia Brain Bank (SWDBB), University of Bristol, UK; Alzheimer’s Disease Research Centre, University of California, Irvine (ADRC-UCI), USA; and National Institute for Health (NIH) NeuroBioBank, USA. This study was approved by the respective research ethics committees. Samples were selected with advice from each brain bank on sample availability for matching age at death, sex, Braak and Braak stage and *APOE* genotype across case types. All samples were de-identified and provided by the brain banks with full research consent.

### Case demographics

In this study, we compared frontal cortex (Brodmann area (BA) 10) of individuals who had DS (DSAD), EOAD (without DS) and healthy ageing euploid individuals (HA). EOAD cases of unknown genetic cause were chosen to match age at death of individuals with DSAD, to remove age discrepancy as a confounding factor when comparing the AD-associated proteome or transcriptome. It is not known if there is a genetic cause of AD in these individuals as sequencing has not been carried out to detect mutations in *APP*, *PSEN1* or *PSEN2*. Although these individuals are not part of a family kindred of autosomal dominant AD, it is possible that they have a *de novo* mutation that is disease-causing. Alternatively, these individuals may have a high polygenic risk score for AD. For this study, these EOAD cases are described as having unknown genetic cause. All cases were matched as far as possible for age at death, sex, Braak and Braak stage, and *APOE* genotype.

### Discovery cohort

A discovery cohort of eight DSAD cases, four HA controls and four EOAD cases were sourced from NBTR (Table 1). No significant difference in age or post-mortem interval (PMI) was found between the three case types. The HA group had one female case and three male cases, the DSAD group had four male and four female cases, and the EOAD group had three female cases and one male case. If not available from NBTR, *APOE* genotype was determined by PCR and Hhal restriction digestion (Table 1). The HA group and EOAD groups had one individual with *APOE* ε3ε4 genotype and three individuals with *APOE* ε3ε3 genotype. The DSAD group had two individuals with *APOE* ε3ε4 genotype, five individuals with *APOE* ε3ε3 genotype and one individual with *APOE* ε2ε3 genotype. All four HA cases were Braak and Braak NFT stage 0. Four DSAD cases were Braak and Braak NFT stage VI, and four cases did not have sufficient material available for complete Braak and Braak staging but had neuropathological examination reports of severe AD pathology, as defined by the presence and severity of amyloid-β plaques and NFT. All four EOAD cases were Braak and Braak NFT stage VI.

**Table 1:** Case demographics for discovery proteomics cohort. Cases (n=4 HA, n=8 DSAD, n=4 EOAD) were selected from Newcastle Brain Tissue Resource and matched where possible for age at death, sex, Braak and Braak stage, and *APOE* genotype. No significant difference in age at death (Univariate ANOVA, F(2,13) = 2.640, p = 0.109) or PMI (Univariate ANOVA, F(2,13) = 0.698, p = 0.515) was found between case types. HA = healthy aging, DSAD = Down syndrome-Alzheimer’s disease, EOAD = early-onset Alzheimer’s disease, APOE = Apolipoprotein E, N/A = not applicable. ‘-‘ = data not available.

| Case type and ID | Braak and Braak stage / Pathology diagnosis | CERAD score | APOE genotype | Sex | Age at AD onset (years) | Age at death (years) | Post-mortem interval (hours) |
| --- | --- | --- | --- | --- | --- | --- | --- |
| HA1 | 0 | - | 33 | M | N/A | 46 | 24 |
| HA2 | 0 | - | 34 | M | N/A | 53 | 19 |
| HA3 | 0 | - | 33 | F | N/A | 59 | 19 |
| HA4 | 0 | - | 33 | M | N/A | 47 | 29 |
| DSAD1 | Down syndrome with Alzheimer's disease pathology | - | 33 | F | 44 | 45 | 9 |
| DSAD2 | Alzheimer's disease with severe plaque and tangle pathology | - | 33 | F | - | 48 | 24 |
| DSAD3 | Down syndrome with severe Alzheimer's disease | - | 34 | F | 52 | 58 | 48 |
| DSAD4 | Down syndrome with Alzheimer's disease pathology | 3 | 33 | M | - | 53 | 84 |
| DSAD5 | VI | 3 | 23 | M | 66 | 67 | 4 |
| DSAD6 | VI | 3 | 34 | F | 57 | 60 | 13 |
| DSAD7 | VI | 3 | 33 | M | 60 | 63 | 40 |
| DSAD8 | VI | 3 | 33 | M | - | 55 | 77 |
| EOAD1 | VI | - | 33 | F | 57 | 67 | 47 |
| EOAD2 | VI | 3 | 33 | M | 55 | 59 | 44 |
| EOAD3 | VI | 3 | 34 | F | 51 | 63 | 11 |
| EOAD4 | VI | 3 | 33 | F | 50 | 58 | 70 |

### Validation Cohort A

A validation cohort of DSAD, HA controls, and EOAD cases (ten cases per group) were sourced from SWDBB (Table 2). This group size was determined by power calculation from experimentally observed effect size and standard deviation for raised APOE abundance observed by mass spectrometry in the discovery cohort (power 80%, α = 0.05). Each group had four male and six female cases. Age at death was significantly higher in HA cases compared to both DSAD and EOAD groups. The HA group had five cases at Braak and Braak NFT stage 0, one case at Braak and Braak NFT stage I and four cases at Braak and Braak NFT stage II. The DSAD group had seven cases of Braak and Braak NFT stage VI, two cases of Braak and Braak NFT stage V and one case of Braak and Braak NFT stage IV. All cases in the EOAD group were Braak and Braak NFT stage VI. As reported by SWDBB, the HA group had six individuals with *APOE* ε3ε3 genotypes, two individuals with *APOE* ε2ε3 genotypes, one individual with *APOE* ε2ε4 genotype and one individual with *APOE* ε3ε4 genotype. The DSAD group had four individuals with *APOE* ε3ε3 genotype, three individuals with *APOE* ε3ε4 genotype, two individuals with *APOE* ε2ε3 genotype, and one individual with *APOE* ε2ε4 genotype. The EOAD group had five individuals with *APOE* ε3ε3 genotype, three individuals with *APOE* ε3ε4 genotype, one individual with *APOE* ε2ε3 genotype, and one individual with *APOE* ε2ε4 genotype.

**Table 2:**
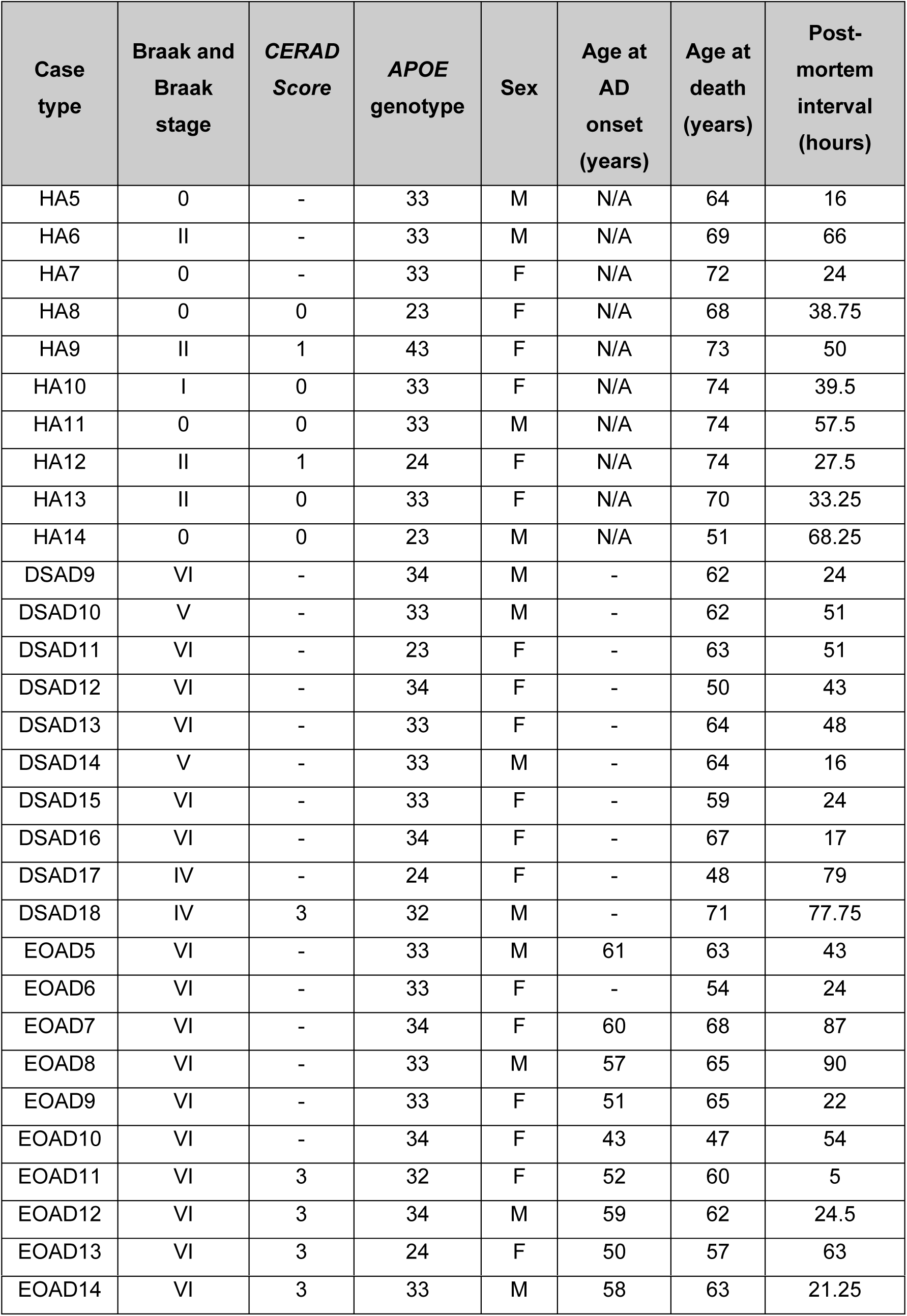
Case demographics for the validation cohort A. Cases were sourced from the South West Dementia Brain Bank (SWDBB). A significant difference in age at death (Univariate ANOVA, F(2,27) = 4.848, p = 0.016) was identified between healthy ageing control (HA) and DSAD (Bonferroni correction, p = 0.046), and HA and EOAD (Bonferroni correction, p = 0.029) cases. No significant difference in PMI (Univariate ANOVA, F(2,27) = 0.008, p = 0.992) was found between case types. HA = healthy aging, DSAD = Down syndrome-Alzheimer’s disease, EOAD = early-onset Alzheimer’s disease, APOE = Apolipoprotein E, N/A = not applicable. ‘-‘ = data not available.

### Validation Cohort B

A validation cohort of DSAD, late-onset AD (LOAD) (n=10 per group), Down syndrome without AD (DS) and age-matched young controls (YC) (n=6 per group) were received from ADRC-UCI and NIH NeuroBioBank (Table 3). Group size for the DSAD and LOAD groups were determined by power calculation from experimentally observed effect size and standard deviation for raised APOE abundance by western blot in validation cohort A (power 80%, α = 0.05). All available frontal cortex samples from individuals below 30 years of age were used for the DS and YC groups. The DSAD and LOAD groups each had five male and five female cases. The DS and YC groups each had five male and one female case. Age at death was significantly different between the groups, with YC and DS groups being significantly younger than both DSAD and LOAD groups, and DSAD also being significantly younger than LOAD (Table 3). All DSAD and LOAD cases were Braak and Braak NFT stage VI, but no Braak and Braak NFT staging was available for YC or DS groups. *APOE* genotype was not available for YC or DS groups. The DSAD group had four individuals with *APOE* ε3ε3 genotype, three individuals with *APOE* ε3ε4 genotype, one individual with *APOE* ε2ε4 genotype, one individual with *APOE* ε2ε3 genotype and one individual with *APOE* ε2ε2 genotype. The LOAD group had four individuals with *APOE* ε3ε3 genotype, five individuals with *APOE* ε3ε4 genotype, and one individual with *APOE* ε2ε4 genotype.

**Table 3:**
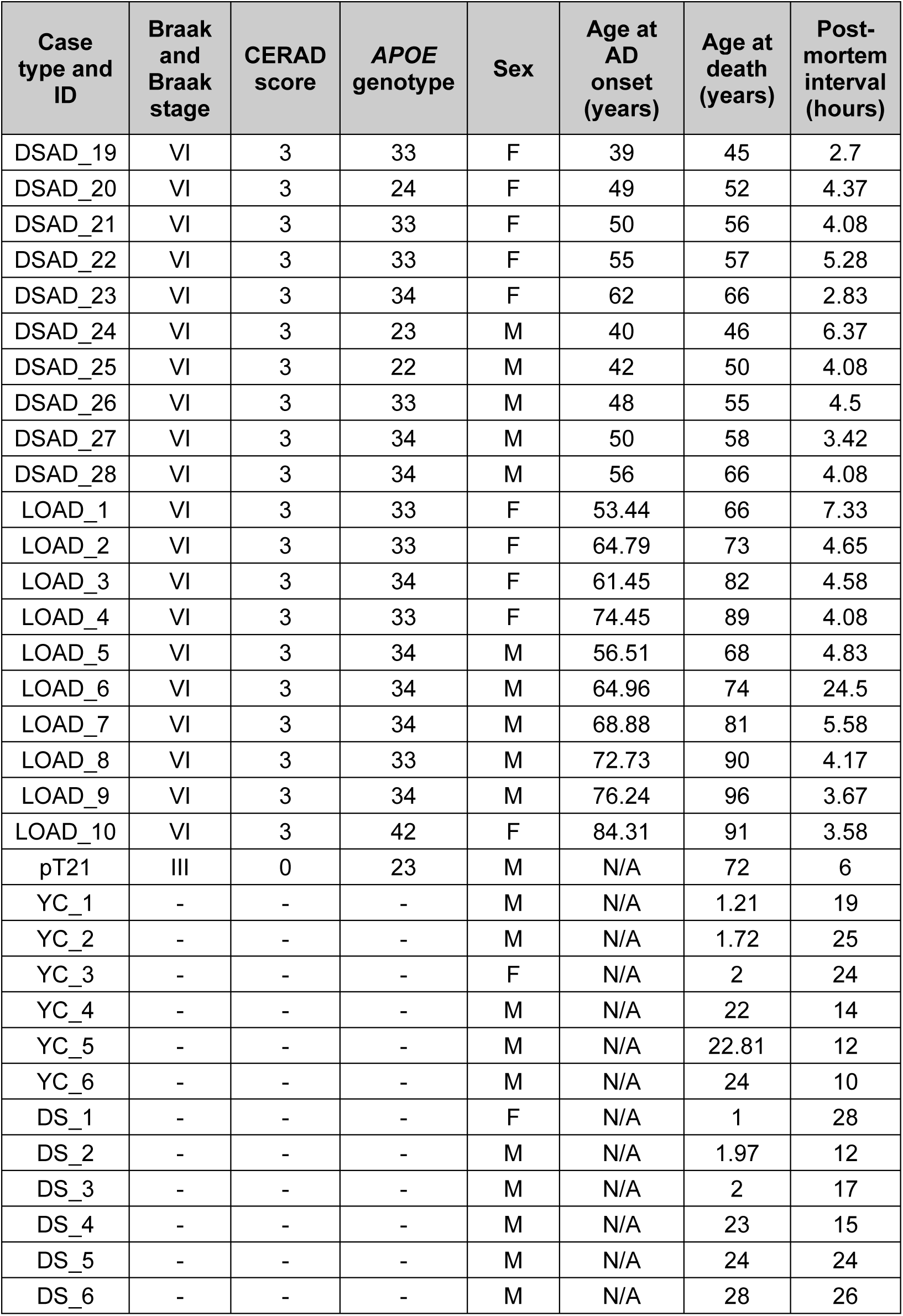
Case demographics for validation cohort B. Cases were sourced from ADRC-UCI and NIH NeuroBioBank (USA). A significant difference in age at death (Univariate ANOVA, F(3,28) = 82.657, p < 0.001) was identified between DSAD and LOAD (Bonferroni correction, p < 0.001), and DSAD and DS (Bonferroni correction, p < 0.001) cases. No significant difference was found between YC and DS cases. A significant difference in PMI (Univariate ANOVA, F(3,28) = 16.683, p < 0.001) was identified between DS cases compared to either DSAD or LOAD cases (Bonferroni correction p < 0.001), and YC cases compared to either DSAD or LOAD (Bonferroni corrections p = 0.003). YC = Young control, DS = Down syndrome, DSAD = Down syndrome-Alzheimer’s disease, LOAD = late-onset Alzheimer’s disease, APOE = Apolipoprotein E, N/A = not applicable. ‘-‘ = data not available.

### APOE Genotyping

DNA was extracted from 20-50 mg of frozen human post-mortem brain material from cases with unknown *APOE* status and positive control samples using the DNeasy Blood and Tissue kit (Qiagen, 69504) according to the manufacturer’s instructions. Briefly, in an MSC Class I hood, chipped tissue was cut into small pieces using a scalpel and mixed with Buffer ATL and proteinase K and incubated at 56 °C, with vortexing, until tissue was completely lysed. Lysed tissue was mixed with Buffer AL and 100 % ethanol and ran through a DNA extraction column. The column was sequentially washed with buffers AW1 and AW2. DNA was eluted in buffer AE. A PCR reaction was carried out on eluted DNA using the HotStarTaq DNA Polymerase kit (Qiagen, 203203) and primers for *APOE*, forward: TCGGCCGCAGGGCGCTGATGG, reverse: CTCGCGGGCCCCGGCCCCGGCCTGGTA. An initial heat activation step was carried out on all samples at 95 °C for 15 min, followed by 18 cycles of denaturation at 95 °C for 30 s, annealing at 70 °C for 1 min, extension at 72 °C for 1 min, and 22 cycles of denaturation at 95 °C for 15 s, annealing at 55 °C for 1 min, extension at 72 °C for 1 min, and a final incubation at 72 °C for 10 min. PCR product was digested using restriction enzyme HhaI (New England Biolabs, R0139S) at 37 °C for 2 h 30 min, followed by 68 °C for 20 min. Digested PCR product of unknown and positive control samples, and O’RangeRuler 10bp DNA Ladder (Thermo Fisher, SN1313), were loaded onto a 5% agarose gel (3 % MetaPhor agarose (Lonza, 50181) and 2 % UltraPure Agarose (Thermo Fisher, 16500500)), ran for 2 h at 135 V, and imaged on a Gel-Doc XR+ Gel Documentation System (Bio-Rad). For all samples, bands were detected at 127, 18 and 16 bp. For an *APOE* ε2 allele, additional bands were detected at 91 and 85 bp. For an *APOE* ε3 allele, additional bands were detected at 91, 48 and 38 bp. For an *APOE* ε4 allele, additional bands were detected at 72, 48 and 38 bp.

### Proteomic mass spectrometry sample preparation

Frozen tissue from discovery cohort cases (Table 1) was homogenised in 50 mM Ambic buffer with 2 % ASB-14 using Precellys 24 homogenizer (Bertin Instruments, P002391-P24T0-A.0) in CK-14 tubes with a 6,500-speed cycle of 2 times 20 s with a 5 s rest period between. Protein concentration (mg/mL) was determined using a spectrophotometer and 0.3 mg protein per sample was used. Samples were spun at 8,500 x g for 10 min at 4 °C. Supernatant was precipitated in ice-cold acetone and the pellet mixed with ice-cold acetone by vortexing to remove metabolites followed by centrifugation at 8,500 x g for 10 min at 4 °C. The pellets were air-dried before being resuspended in 70 % formic acid, by shaking overnight at 4 °C. The pellets were then dried by speed-vac for 3 h, reconstituted in 100mM Tris (pH 7.8) with 6M urea, 1% ASB-14 by shaking for 1 h, then reduced by the addition of dithioerythritol (Sigma, 646563) with shaking for 1 h, before the addition of iodoacetamide (Sigma, I1145) with shaking for 30 min. To digest the proteins into peptides, Trypsin/LysC Mix, Mass spec grade enzyme (Promega, V5073) in trypsin buffer was used, at 37 °C overnight. Samples were fractionated on Isolute C18 columns (Biotage, 220-0010-A). Columns were washed with 100 % acetonitrile (ACN) and primed with 20 mM NH_4_OH. Pellet and supernatant fractions were pooled for each sample and loaded onto the columns’, adhered proteins were washed with 20 mM NH_4_OH and eluted with increasing concentrations of ACN (7.4 %, 14 %, 20.4 %, 60 %) generating four fractions. ACN was removed by speed vacuum for 6 h and pellets were stored at -20 °C.

### Quantitative label-free mass spectrometry and analysis

Label free proteomics analysis was performed as previously described [8]. Briefly, fractionated peptides were reconstituted with 5 % ACN 0.1 % TFA. 0.3 µg of peptides was injected onto a NanoAquity and separated by reverse phase chromatography over a 60 min gradient. Peptides were detected using a SYNAPT G2-Si High-Definition mass spectrometer. Case EOAD4 was excluded from discovery analysis due to poor chromatographic separation of peptides, however, it was included in all subsequent studies. All raw data was loaded into Progenesis QI for proteomics software and processed with the following settings: low energy threshold intensity was 200 counts, and elevated energy threshold was 20 counts; elution start set to 10 min and elution finish set to 50 min, manual alignment was used. Peptides were identified using the UniProt Human Reference Proteome database 2022, search parameters were as follows: trypsin digest reagent; max 3 missed cleavages; max protein mass 800 kDa, post-translational modifications Carbamidomethyl C, oxidation M. Tolerance parameters had a false-discovery rate less than 1 %. Ion matching requirements were set as follows: 2 or more fragments/peptides, 3 or more fragments/proteins, 1 or more peptides/protein. Proteins with <2 unique peptides were excluded. All significantly altered proteins were ranked by fold-change and a 1.25-fold-change cut off for both up and down-regulated proteins was applied to examine alterations between DSAD, EOAD and HA controls. The mass spectrometry proteomics data have been deposited to the ProteomeXchange Consortium via the PRIDE [49] partner repository with the dataset identifier PXD058779 and 10.6019/PXD058779.

### Single-nuclei RNA sequencing

Nuclei were isolated from frozen frontal cortex from discovery cohort cases (Table 1). Tissue was transferred directly onto ice-cold sucrose buffer and homogenised by hand. The homogenate was layered on top of a sucrose cushion and nuclei were recovered after centrifugation. Nuclei were resuspended in PBS containing BSA (to prevent clumping) and RNase inhibitors, counted and diluted to 1,000 nuclei/μl. Sequencing libraries were prepared using the “single cell 3’ library kit v3” with the Chromium instrument (10X Genomics) aiming to obtain libraries for ∼5,000 independent nuclei per sample. Libraries were sequenced on an Illumina NovaSeq6000 instrument (UCL Genomics), to an average of at least 100,000 reads/nucleus.

Raw sequencing data were processed through the software CellRanger (10X Genomics) to assign each transcript read to its respective nucleus, we performed alignment to the transcriptome, counting reads, and processing of the unique molecular identifier tag to reduce PCR bias [72]. The resulting gene count table was analysed using the Seurat package (version 5) [24]. Quality control was performed to remove low-quality cells based on the number of detected genes and unique molecular identifiers (UMIs). Cells with an unusually high number of UMIs, suggesting potential doublet events, were also excluded using scDblFinder [18]. Using SCTransform, gene expression data were normalised to adjust for differences in sequencing depth between cells and scaled to correct for cell-to-cell variation in total UMI count [22]. Data were also normalised to remove confounding sources of variation such as mitochondrial mapping percentage, post-mortem delay, age of death, and batch number.

Principal component analysis (PCA) was performed on the scaled data to reduce the dimensionality of the dataset. The top principal components explaining the most significant variance in the data were retained for downstream analysis. The PCA-generated principal components were used for clustering cells using the graph-based Louvain algorithm implemented in Seurat. Cluster annotation was performed to identify putative cell types based on known marker genes. Marker genes for various cell types were obtained from the literature and publicly available databases [35]. Cell types were annotated by comparing the expression pattern of marker genes within each cluster.

The snRNAseq data was visualised using a Uniform Manifold Approximation and Projection (UMAP) to view the cell clusters in two dimensions. Dot plots were generated to visualise the expression patterns of marker genes and differentially expressed genes (DEG) in each cell type. DEG analysis was performed to identify genes that were significantly differentially expressed between identified cell clusters. This analysis was carried out using the "FindMarkers" function in Seurat, employing Wilcoxon rank-sum analyses. Any clusters with less than four nuclei were omitted from DEG analysis. All statistical analyses were performed using R (version 4.2.1) and relevant packages, as specified in the text. Statistical significance was defined as a false discovery rate of < 0.05. The data discussed in this publication have been deposited in NCBI’s Gene Expression Omnibus [13] and are accessible through GEO Series accession number GSE284141.

Data from the published snRNAseq dataset [46] was re-analysed to examine the expression of key target genes between DSAD and healthy control cases used in that study. The data was downloaded, and Scanpy toolkit in Python was used for log transformation and normalizing the raw data. Violin plots were used to visualize the expression of Hsa21 and non-Hsa21 associated genes in DSAD versus healthy control samples. Mann-Whitney U non-parametric test was employed to determine the statistical significance of the observed differences.

### RNA preparation, RNA integrity assay and quantitative PCR

RNA was extracted using the Monarch Total RNA Miniprep Kit (NEB, T2010) according to the manufacturer’s instructions using Proteinase K digestion at 55 °C for 5 min. Genomic DNA was removed by gDNA column, prior to application to an RNA Purification Column for on-column DNase treatment (DNase I for 15 min at room temperature), prior to washing and elution in RNase-free water. RNA Integrity number (RIN) was measured using an RNA ScreenTape Assay on the Agilent 4200 TapeStation system (Agilent, G2991AA) using RNA buffer (Agilent, 5067-5577). cDNA was produced using the SuperScript™ First-Strand Synthesis System for RT-qPCR kit (Thermo Fisher, 11904-018) according to the manufacturer’s instructions. Samples were incubated at 65 °C for 5 min with 10 mM dNTP mix and Random hexamers, prior to the addition of 10X RT Buffer, 25 mM MgCl2, 0.1 M DTT and RNaseOUT and SuperScript™ II RT enzyme, for the production of cDNA; room temperature for 10 mins, 42 °C for 50 min, 70 °C for 15 min. RNA was then digested with RNase H at 37°C for 20 min. Taqman RT-qPCR system was used to perform RT-qPCR, using TaqMan™ Multiplex Master Mix (Thermo Fisher, 4461881), *ACTB* TaqMan Assay (Reported dye: VIC) (Hs0302943_g1), *GAPDH* TaqMan Assay (Reporter dye: JUN) (Thermo Fisher, 4486527) and *APOE* TaqMan Assay (Hs00171168_m1, Reporter dye: FAM). Samples were run on fast mode on a Quantstudio 3 Real-time PCR system (Thermo Fisher, A28137), to generate a cycle threshold (Ct) for the housekeeping genes and genes of interest. An average of the triplicate Ct values per sample was used. Relative gene expression was calculated using the 2^−ΔΔCt^ method.

### Western Blot

50-100 mg of frozen tissue was homogenised in 500 µL T-PER Tissue Protein Extraction Reagent (Thermo Fisher, 78510) with PhosSTOP phosphatase inhibitor (Roche, 04906837001) and cOmplete Protease Inhibitor Cocktail (Roche, 04693116001), using a Tissue Ruptor with disposable probes (Qiagen, 990890). The lysate was centrifuged at 10,000 x g for 30 min at 4 °C. Supernatant was aliquoted and stored at -70 °C. Total protein concentration was calculated using Protein Assay Dye Reagent (Bio-Rad, 5000006) in triplicate and read at 595 nm using a microplate reader (Tecan Spark, ZT2973521S), or by Pierce BCA Protein Assay kit (Thermo-Fisher, 23225).1 µg/µL of protein from each sample was prepared in NuPAGE™ LDS Sample Buffer (4X) (Thermo Fisher, NP0007), and NuPAGE™ Sample Reducing Agent (10X) (Thermo Fisher, NP0004) denaturing at 95 °C for 5 min. Linearity of signal was experimentally determined. 5-15 µg protein was loaded per lane of 26-well NuPAGE™ 4 to 12 %, Bis-Tris, 1.0 mm, Midi Protein Gels (Thermo Fisher, WG1403BOX) or to a 26-well, 18-well, or 12+2-well, 4-12% Criterion XT Bis-Tris Protein Gel (Bio-Rad, 3450125) with Chameleon Duo Pre-stained Protein Ladder (Licor Bio, 928-60000). Sample loading order was randomised. Gels were run in 1X NuPAGE™ MES SDS Running Buffer (Thermo Fisher, NP0002) at 100 V, 400mA and 120 Watts for 120-180 min. Proteins were transferred to PVDF membranes using the Trans-Blot Turbo Midi 0.2 µm PVDF Transfer Packs (Bio-Rad, 1704157) in the Transblot Turbo Transfer System (Bio-Rad, 1704150) at 25 V and 2.5 Amp for 7-20 min. Membranes were stained for total protein using the Revert 700 Total Protein Quantification kit (Licor Bio, 926-11016). After total protein quantitation and washing, membranes were blocked for 1 h in Intercept® Blocking Buffer (Licor Bio, 927-70001) and incubated in primary antibodies (APOE (Calbiochem, 178479), APOE (Sigma, SAB2701949), APP (Y188, abcam, Ab220793), S100B (Abcam, Ab41548), Tau (Dako, A0024), AT8 (Thermo Fisher, MN1020)) diluted in blocking buffer overnight at 4 °C. Followed by incubation with secondary antibody (IRDye® 680RD Goat anti-Mouse IgG Secondary Antibody (Licor Bio, 926-68070), IRDye® 800CW Goat anti-Rabbit IgG Secondary Antibody (Licor Bio, 926-32211) or IRDye® 800CW Donkey anti-Goat igG Secondary Antibody (Licor Bio, 926-32214) diluted in blocking buffer for 1 h at room temperature in the dark with rocking, prior to imaging using the Odyssey® CLx Imaging system (Licor Bio). For re-probing, membranes were stripped for 20 min with rocking using Restore™ Western Blot Stripping Buffer (Thermo Fisher, 21059), blocked for 1 h and re-incubated with antibody. Empirica Studio v3.2 (Licor Bio) was used to quantify protein bands and normalise to total protein in the same lane. All samples were normalised against the average HA or YC control value within a blot. A minimum of two technical replicates per sample were then averaged.

### Protein fractionation and amyloid-β quantification

Fractionation was carried out as previously described [58]. Total brain sample was weighed prior to homogenisation in three-volumes of TBS (50mM TBS-HCL (Fisher Scientific, 10776834), pH8) containing PhosSTOP phosphatase and cOmplete Protease Inhibitor Cocktails, with a Tissue Ruptor and disposable probes. Samples were then balanced prior to centrifugation at 186,000 x g in an Optima Max-XP Benchtop Ultracentrifuge (Beckman Coulter, 393315) fitted with a fixed-angle rotor TLA-55 (Beckman Coulter, 366725) for 30 min at 4 °C. The supernatant is the TBS fraction and was aliquoted, snap frozen and stored at - 70 °C (soluble fraction). The pellet was resuspended in three-volumes of original tissue weight in ice-cold Triton-X-100 buffer (1 % Triton-X-100 (Sigma, X100-5ML) in TBS, pH8) containing protease and phosphatase inhibitors, balanced, and centrifuged at 186,000 x g for 30 min at 4 °C. Resulting Triton supernatant was aliquoted, snap frozen and stored at -70 °C (membrane-associated fraction). The pellet was resuspended in three-volumes of Guanidine-HCl buffer (5M guanidine-HCl (Thermo Fisher, 24110) in TBS, pH 8) containing protease and phosphatase inhibitors, incubated overnight with shaking at 4 °C, aliquoted and stored at - 70 °C (plaque-associated/aggregated fraction).

Amyloid-β_38_, _40_ and _42_ isoforms were quantified in fractionated protein samples using the MSD V-PLEX Aβ Peptide Panel 1 (6E10) Kit (MesoScale Discovery, K15200E-2). The Tris and Triton fractions were diluted 1:4, and the Guanidine fraction was diluted 1:10,000 in kit provided Diluent 35. Analyte binding was quantified by a SULFO-TAG conjugated anti-Aβ 6E10 detection antibody, using a MSD SECTOR S 600 plate reader. Discovery Workbench 4.0 software was used to analyse plate data based on the standard curve. Analyte concentration (pg/mL) was normalised to tissue weight per buffer volume homogenised in to give a final concentration (pg/mg) which was used for statistical analysis.

### Experimental design and statistical analysis

Individual post-mortem brain donor was used as the unit of replication. Unique identification numbers were assigned to all samples to ensure the study remained blind and for technical randomisation during data acquisition and analysis. Statistical analysis was carried out using SPSS Statistics 29 (IBM) for ANOVA and Prism 10 (GraphPad) for correlation analysis. When ANOVA was used for analysis, sex and case type (HA, EOAD or DSAD) were used as variables, and age at death and post-mortem interval (PMI) as covariates. We also repeated our analysis including *APOE* genotype as a variable for the analysis of APOE abundance by western blot and mass spectrometry but note this is exploratory as we have a low number of cases, particularly of *APOE* ɛ2ɛ3 and ɛ2ɛ4 alleles, in this case series. Graphs were plotted using Prism 10 (GraphPad), data are presented as mean ± standard error of the mean (SEM), and p-values < 0.05 were considered to be statistically significant.

## Results

### Label-free proteomic comparison of DSAD, EOAD and healthy ageing identifies proteins that have changed abundance resultant from trisomy 21

To understand how an extra copy of Hsa21 changes the abundance of proteins within the brain in the context of AD, we quantified the abundance of proteins by label-free mass spectrometry of the frozen frontal cortex (BA10) in individuals with DSAD, EOAD and HA controls. We identified 2855 proteins with 2 or more unique peptides in the study; we observed that 362 proteins were significantly different between case types (Supplementary Information 1; Table 4-6, Figure 1).

**Figure 1.**
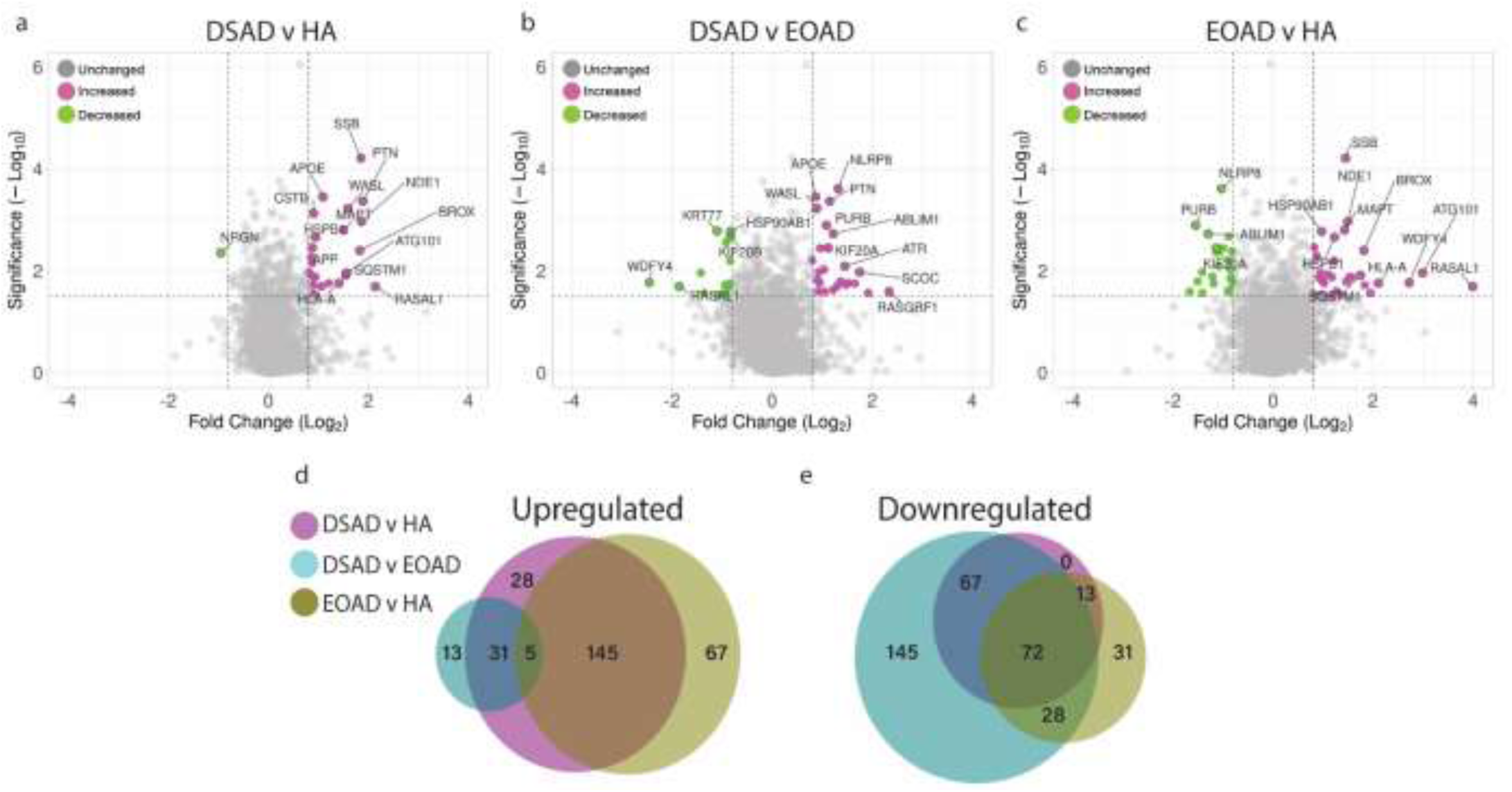
Significantly up- and down-regulated proteins between DSAD, EOAD and HA control frontal cortex by label-free proteomics. Volcano plots show Log_2_(Fold-change) between case types, (a) DSAD compared with HA control, (b) DSAD compared with EOAD, (c) EOAD compared with HA control, plotted against - Log_10_(ANOVA p), using fold-change threshold of ±0.8 and a significance threshold of 1.5, created using VolcaNoseR [19]. The top 15 dysregulated proteins are labelled on each plot. Venn diagrams, created in BioVenn [29], demonstrate the common (d) upregulated and (e) downregulated proteins across the three comparisons of DSAD v HA, DSAD v EOAD and EOAD v HA.

**Table 4:** Significantly increased chromosome 21 encoded proteins in DSAD frontal cortex. Significantly altered Hsa21-encoded proteins between case types (ANOVA, p < 0.05), which have increased abundance (>1.25-fold) in DSAD frontal cortex compared with both HA and EAOD matched controls. Magenta represents upregulated proteins, green represents downregulated proteins, and greater intensity of colour indicates a greater degree of change. HA = healthy aging, DSAD = Down syndrome-Alzheimer’s disease, EOAD = early-onset Alzheimer’s disease.

| Protein name | Gene name | Case type ANOVA (p) | Fold change |  |  |
| --- | --- | --- | --- | --- | --- |
|  |  |  | DSAD versus HA | DSAD versus EOAD | EOAD versus HA |
| Pyridoxal kinase | *PDXK* | 8.78E-07 | 1.54 | 1.61 | -1.04 |
| Cystatin B | *CSTB* | 0.0007416 | 1.87 | 1.72 | 1.09 |
| Synaptojanin-1 | *SYNJ1* | 0.0031671 | 1.20 | 1.30 | -1.08 |
| Amyloid precursor protein | *APP* | 0.0035328 | 1.84 | 1.12 | 1.64 |
| Protein S100-B | *S100B* | 0.0070407 | 1.83 | 1.39 | 1.32 |
| Carbonyl reductase [NADPH] 1 | *CBR1* | 0.0107494 | 1.76 | 1.29 | 1.37 |
| Phosphofructokinase liver type | *PFKL* | 0.0120288 | 1.36 | 1.67 | -1.23 |
| Carbonyl reductase [NADPH] 3 | *CBR3* | 0.0397656 | 1.23 | 1.04 | 1.19 |

In this study, we primarily aimed to identify proteins that had altered abundance due to trisomy 21. Thus, we first focused on Hsa21-encoded proteins within the dataset which we hypothesised would have increased abundance due to the dosage imbalance caused by the extra copy of Hsa21. We identified 23 Hsa21-encoded proteins (Supplementary Information 1) and found that 8 of these were significantly increased in DSAD cases compared with either HA or cases of EOAD (Table 4). These findings are consistent with previous reports of increased abundance of APP (amyloid precursor protein), S100B (S100 calcium-binding protein B), and SYNJ1 (synaptojanin-1) in the frontal cortex and CSTB (cystatin-B) in the temporal cortex of people with DSAD [42, 55, 69]. Our finding of raised PDXK (pyridoxal kinase), contrasts with a previous report that the protein was not elevated in the brains of individuals with DS at ∼ 19 weeks gestation [59]. Raised abundance of CBR1 (carbonyl reductase 1), CBR3 (carbonyl reductase 3) and PFKL (6-phosphofructokinase) in the brain of individuals with DSAD has not previously been reported. We notably detected multiple Hsa21-encoded proteins, such as CCT8 and USP16, in the frontal cortex with similar abundance in all three case types (CCT8 HA x̄ = 62003, DSAD x̄ = 64142, EOAD x̄ = 62610, and USP16 HA x̄ = 868, DSAD x̄ = 873, EOAD x̄ = 844) (Supplementary Information 1). Thus, an additional copy of a Hsa21 gene due to trisomy does not always result in an elevation of protein abundance.

To identify proteins encoded by genes on other chromosomes with altered abundance in DSAD compared to EOAD and HA controls, we examined all significantly changed proteins (Supplementary Information 1) that were altered ±1.25-fold. This identified 31 robustly upregulated proteins that were not encoded by Hsa21 (Table 5, Figure 1 a-d). SSB (Lupus La protein), encoded on chromosome 2, is an RNA binding protein with key roles in the regulation of transcription, translation, and splicing, and was the most significantly altered protein upregulated in DSAD compared to both EOAD and HA (Figure 1 a, c). The abundance of PTN (pleiotrophin), an injury response cytokine secreted from microglia, macrophages, astrocytes, and endothelial cells in the brain [14, 63, 70], encoded on chromosome 7, was also increased in DSAD (Table 5, Figure 1 a, b). WASL (Neural Wiskott-Aldrich syndrome protein), encoded by chromosome 7, involved in actin cytoskeleton reorganisation in the brain [34], and NDE1 (Nuclear distribution protein nudE homolog 1), a centromere regulatory protein encoded by chromosome 16, were both also found to have increased abundance in the DSAD cases (Table 5, Figure 1 a, b). Notably, we identified that APOE (apolipoprotein E), encoded on chromosome 19, was upregulated in DSAD compared to EOAD and HA controls (Table 5, Figure 1 a, b). APOE is a lipoprotein involved in the transport of cholesterol and phospholipids between cells, and allelic variation in the *APOE* gene is a strong genetic risk factor for AD [16, 41]. We identified 64 unique peptides that mapped to APOE, and 16 of these were significantly elevated in DSAD cases, suggesting upregulation of the proteins’ abundance in the frontal cortex is caused by trisomy of Hsa21 (Supplementary Information 1).

**Table 5:**
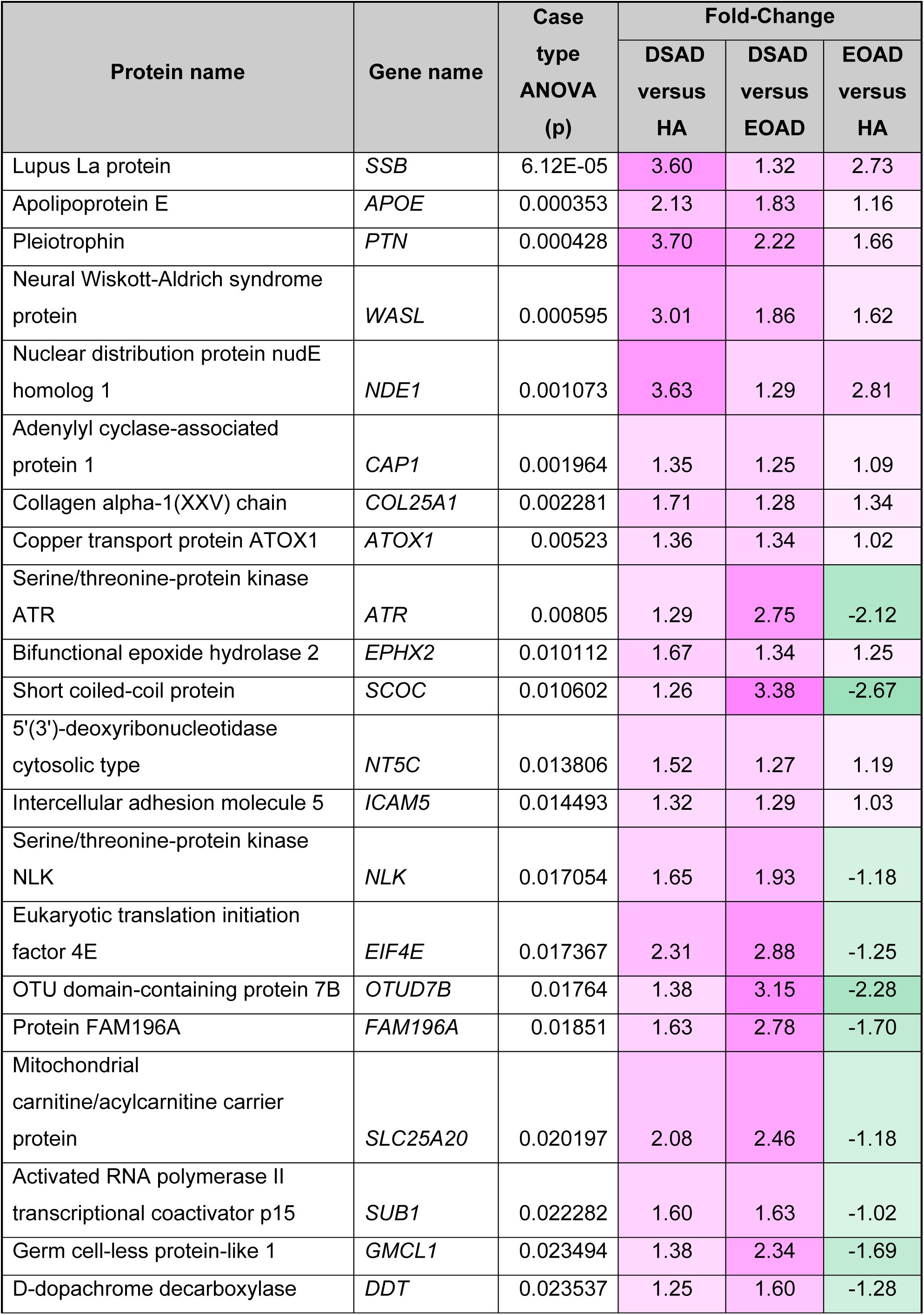
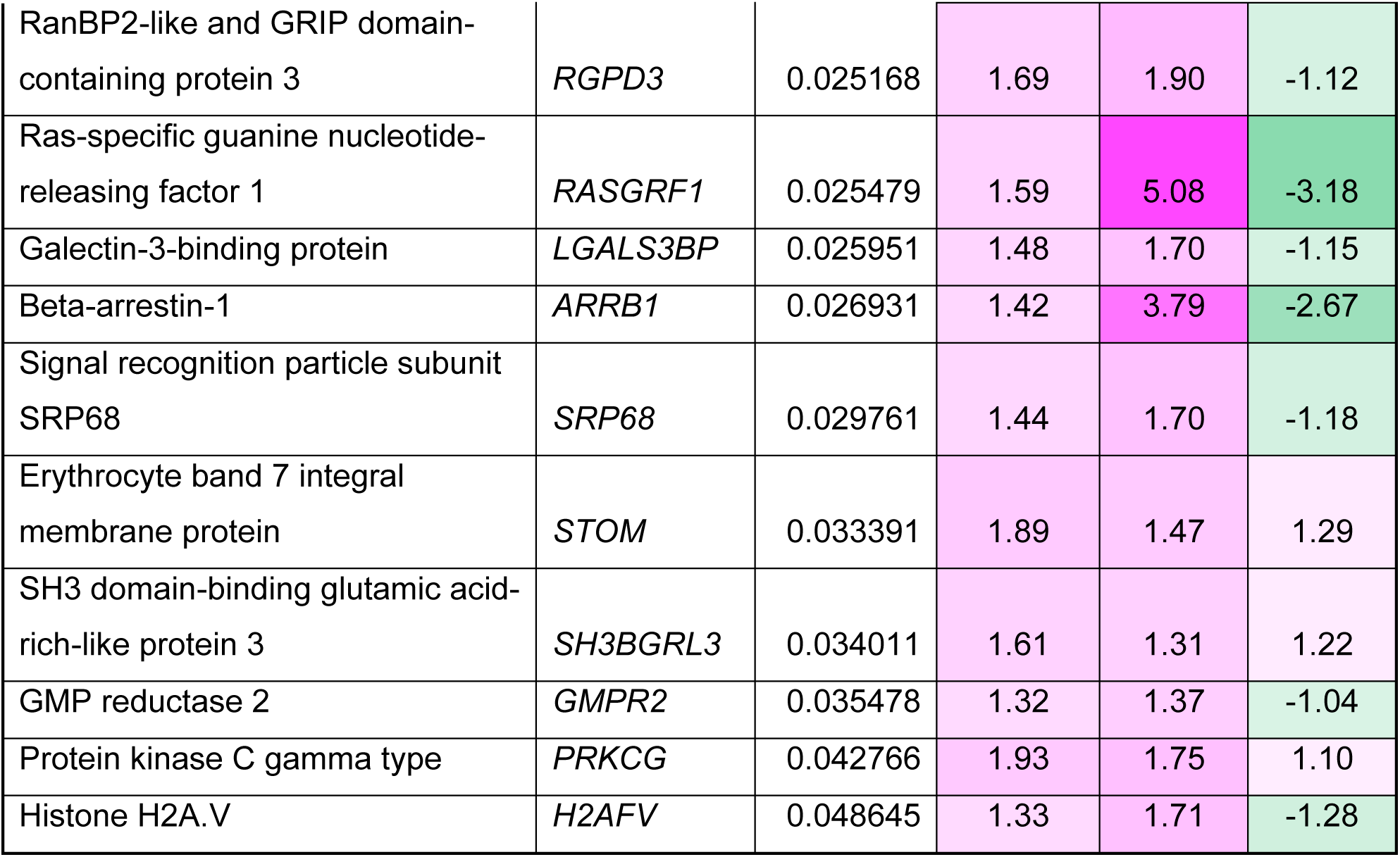
Significantly increased non-chromosome 21 encoded proteins in DSAD frontal cortex. Significantly altered proteins, not encoded by Hsa21, between case types (ANOVA, p < 0.05), which have increased abundance (>1.25-fold) in DSAD frontal cortex compared with both HA and EOAD matched controls. Magenta represents upregulated proteins, green represents downregulated proteins, and greater intensity of colour indicates greater degree of change. HA = healthy aging, DSAD = Down syndrome-Alzheimer’s disease, EOAD = early-onset Alzheimer’s disease.

We also identified 19 proteins encoded by chromosomes other than Hsa21 that were downregulated in DSAD compared to both EOAD and HA controls (Table 6). This included the novel identification of KRT77 (keratin 77), KIF20B (Kinesin family member 20B), UQCRB (Cytochrome b-c1 complex subunit 7), INPP4A (Type I inositol 3-4-bisphosphate 4-phosphatase) and ALDH1A1 (Aldehyde Dehydrogenase 1 Family Member A1) that were downregulated in the brains of individuals with DSAD compared with control groups (Table 6, Figure 1). 145 proteins were commonly upregulated, and 45 proteins were commonly downregulated, in DSAD v HA and EOAD v HA, suggesting these proteins were dysregulated due to similar mechanisms of AD (Figure 1 d, e).

**Table 6:** Significantly decreased non-chromosome 21 encoded proteins in DSAD frontal cortex. Significantly altered proteins, not encoded by Hsa21, between case types (ANOVA, p < 0.05), which have decreased abundance (>1.25-fold) in DSAD frontal cortex compared with both HA and EOAD matched controls. Magenta represents upregulated proteins, green represents downregulated proteins, and greater intensity of colour indicates a greater degree of change. HA = healthy aging, DSAD = Down syndrome-Alzheimer’s disease, EOAD = early-onset Alzheimer’s disease.

| Protein name | Gene name | Case type ANOVA (p) | Fold-Change |  |  |
| --- | --- | --- | --- | --- | --- |
|  |  |  | DSAD versus HA | DSAD versus EOAD | EOAD versus HA |
| Keratin_ type II cytoskeletal 1b | *KRT77* | 0.00164 | -1.35 | -2.14 | 1.58 |
| Kinesin-like protein KIF20B | *KIF20B* | 0.002076 | -1.40 | -1.79 | 1.27 |
| Cytochrome b-c1 complex subunit 7 | *UQCRB* | 0.002157 | -1.40 | -1.26 | -1.11 |
| Type I inositol 3_4-bisphosphate 4-phosphatase | *INPP4A* | 0.002799 | -1.25 | -1.89 | 1.52 |
| Aldehyde Dehydrogenase 1 Family Member A1 | *ALDH1A1* | 0.006048 | -1.31 | -1.26 | -1.04 |
| Syntaxin-1B | *STX1B* | 0.010062 | -1.28 | -1.30 | 1.02 |
| ESF1 homolog | *ESF1* | 0.012203 | -1.57 | -1.26 | -1.24 |
| Programmed cell death protein 10 | *PDCD10* | 0.01242 | -1.45 | -1.66 | 1.15 |
| Torsin-3A | *TOR3A* | 0.013997 | -1.37 | -1.47 | 1.07 |
| WASH complex subunit 4 | *WASHC4* | 0.017995 | -1.46 | -1.80 | 1.24 |
| Synaptotagmin-17 | *SYT17* | 0.020512 | -1.46 | -1.62 | 1.11 |
| NK-tumor recognition protein | *NKTR* | 0.02079 | -1.26 | -1.49 | 1.18 |
| Vascular endothelial growth factor receptor 1 | *FLT1* | 0.020816 | -1.37 | -1.86 | 1.36 |
| Pyruvate Dehydrogenase Phosphatase Catalytic Subunit 1 | *PDP1* | 0.033151 | -1.37 | -1.48 | 1.08 |
| Ubiquinol-Cytochrome C Reductase Hinge Protein | *UQCRH* | 0.038237 | -1.34 | -2.83 | 2.11 |
| 40S ribosomal protein S5 | *RPS5* | 0.041307 | -1.26 | -1.37 | 1.09 |
| ADP/ATP translocase 2 | *SLC25A5* | 0.044547 | -1.25 | -1.32 | 1.06 |
| NADH dehydrogenase [ubiquinone] 1 beta subcomplex subunit 7 | *NDUFB7* | 0.046316 | -1.29 | -1.92 | 1.49 |
| Antithrombin-III | *SERPINC1* | 0.047911 | -1.38 | -1.78 | 1.29 |

### The transcriptome is altered in the frontal cortex of individuals with DSAD compared with EOAD and HA in a range of cell types

To determine which cell types may contribute to the identified changes in protein abundance in DSAD, we undertook a snRNAseq study in the same cases which underwent proteomic analysis (discovery cohort, Table 1). We sequenced the transcriptome of nuclei gathered from the 16 cases. After quality control preprocessing, we recovered a total of 89,649 nuclei across all samples of which we identified a total of 38 molecularly unique clusters (Figure 2 a). Less than 5000 nuclei were obtained from four cases; two DSAD cases (DSAD6, DSAD8) and two EOAD cases (EOAD2, EOAD3), but we have included these cases in our analysis to maintain a balanced sample size (Figure 2 b, Supplementary Information 2). RNA integrity was not found to correlate with the number of nuclei recovered (Pearson’s R = 0.2255 p = 0.4382) (Supplementary Information 2). We noted that for multiple cell types less than 10 nuclei were recovered from some cases, thus any identified differential expression should be treated as preliminary and requires independent validation (Figure 2 a, b, Supplementary Information 2).

**Figure 2.**
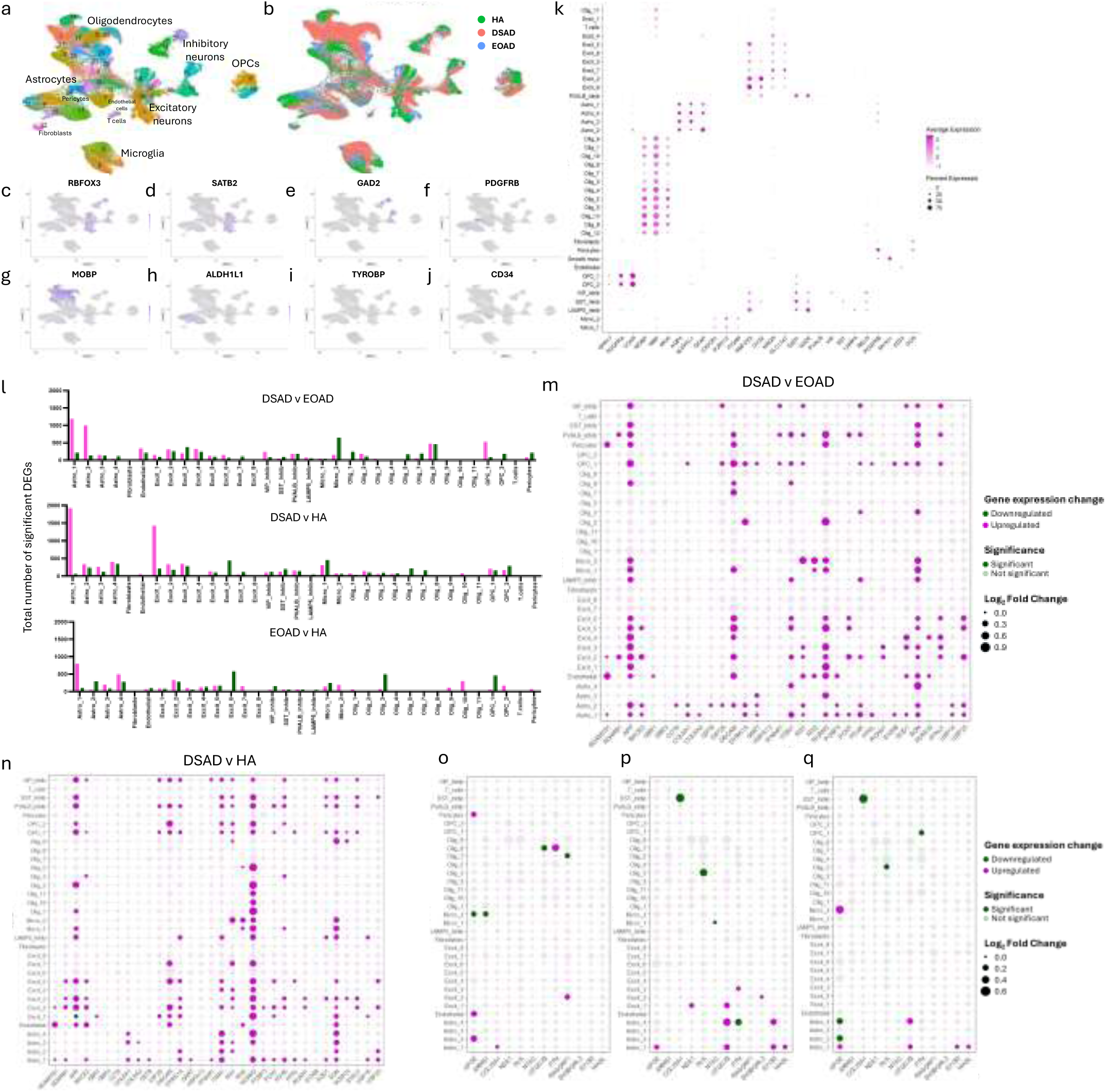
Single-nuclei RNA sequencing demonstrates differential expression of chromosome 21 genes in a broad range of cell types, and upregulation of *APOE* in astrocytes, endothelial cells, and pericytes, in DSAD compared with EOAD and HA controls. (a) Annotated UMAP demonstrates nuclei clusters identified. (b) UMAP demonstrates the identified nuclei by case type. (c-j) UMAP demonstrates the molecular identity of each cluster; neurons (*RBFOX3*), excitatory neurons (*SATB2*), inhibitory neurons (*GAD2*), pericytes (*PDGFRB*), oligodendrocytes (*MOBP*), astrocytes (*ALDH1L1*), microglia (*TYROBP*), and endothelial cells (*CD34*). This is further elaborated on in (k) with a dot plot showing multiple cellular markers used to identify cell clusters. (l) The total number of differentially expressed genes (DEGs) detected across nuclei clusters (magenta = upregulated, green = downregulated). (m, n) Hsa21-encoded genes identified in the proteomics dataset, and other commonly investigated Hsa21 genes, are significantly upregulated across multiple cell types in DSAD compared to EOAD and HA, respectively. (o-p) Non-Hsa21 genes which were significantly different between case types in the proteomic study are represented in dot plots. (o, p) *APOE* is significantly upregulated in astrocytes in DSAD compared to HA, and in astrocytes, endothelial cells and pericytes in DSAD compared to EOAD.

Of the 38 clusters, we identified all the main neural cell types (Figure 2 c-k). This included eight excitatory neuronal clusters, four distinct inhibitory neuronal clusters (*VIP*, *SST*, *PVALB*, *LAMP5*), as well as glial cells; 13 oligodendrocyte, two microglial, four astrocytic, and two oligodendrocyte precursor cell (OPCs) clusters. We also collected nuclei expressing endothelial cell, T cell, fibroblast, and pericyte markers. Differential gene expression identified increased transcriptomic variability in the astrocytic and excitatory neuron clusters of DSAD when compared to EOAD and HA groups, with fewer but significant changes in gene expression seen in microglia, isolated oligodendrocytes, and OPCs clusters (Figure 2 l).

We found increased expression of the Hsa21 genes that also had elevated protein levels in our proteomic study, as well as other Hsa21 genes that we did not identify by mass spectrometry in our proteomic analysis, in DSAD cases compared with euploid controls (Figure 2 m, n). We observed an elevation of Hsa21 gene expression in a range of cell types in DSAD cases compared to HA or EOAD controls. Consistent with a previous report, we found elevated expression of *APP* in many cell types in the frontal cortex (Figure 2 m, n) [48]. Re-analysis of a previously published snRNAseq dataset, also showed the significant upregulation in *APP* expression in astrocytes and microglia in DSAD compared to healthy controls [42]. Together these data highlight that further research on the effect of elevation of this key AD protein in the development of DSAD in cells other than neurons is warranted. Of the Hsa21 proteins we identified to have increased abundance in our proteomic study, *CSTB* was found to have elevated expression in subtypes of astrocytes, and *PDKX* was raised in a broad range of cell types (Figure 2 m, n). *SYNJ1* had increased expression in multiple excitatory and inhibitory neuronal populations, as well as populations of oligodendrocyte precursor cells (OPCs). *CBR1* expression was found to be elevated by trisomy of Hsa21 in a subtype of astrocytes and excitatory neurons (Figure 2 m, n). *S100B* had increased expression in astrocytes and OPCs, and *PFKL* in a subtype of astrocytes, OPCs and excitatory neurons (Figure 2 m, n). In contrast with the proteomic results, *CBR3* did not have significantly increased expression in DSAD brain compared to either HA or EOAD. Although CCT8 and USP16 were not observed to have changed abundance at the protein level (Supplementary Information 1), expression of these Hsa21 genes were elevated in several cell types in the cases of DSAD, demonstrating that altered transcript abundance does not always result in changes to protein levels due to endogenous regulatory mechanisms (Figure 2 m, n). These data are consistent with a previous report comparing bulk proteomics and bulk RNAseq in brain samples from young individuals with DS compared with euploid-matched controls, which also reported a discordance between transcript and protein levels [53]. Moreover, the protein products of some genes that were significantly increased in expression in DSAD in a broad range of cell types, such as *NCAM2* and *SON*, were not identified by mass spectrometry, showing discordance between snRNAseq and proteomic results. This may be affected by the sensitivity and depth of coverage of these experiments, and further validation is required to study targets not identified within these studies. Few Hsa21 genes have significantly altered expression between EOAD and HA in both the proteomic and transcriptomic datasets, showing the specific effect of three-copies of Hsa21 on the expression of genes on this chromosome (Supplementary Figure 1, Table 4).

To understand the cell types that may be contributing to the observed wider dysregulation of the proteome we observed in DSAD, we similarly analysed the cell types in which non-Hsa21 proteins had altered gene expression (Figure 2 o-q). This analysis identified that *APOE* expression was upregulated in endothelial (*CD34* expressing, Figure 2 j), pericytic (*PDGFRB* expressing, Figure 2 f), and astrocytic (*AQP4, ALDH1L* or *GFAP* expressing, Figure 2 h) nuclei in DSAD compared to EOAD, and in a second subtype of astrocytes in DSAD compared to HA (Figure 2 n). In support of this data, re-analysis of a previously published snRNAseq dataset showed the upregulated expression of *APOE* in astrocytes in DSAD compared to healthy controls [46]. Thus, these cells may be the source of the increased abundance of the protein we observed in the frontal cortex of individuals who had DSAD. Moreover, we observed a decrease in the expression of *APOE* in a subtype of microglia (*CX3CR1, ITGAM, AIF1*, or *P2RY12* expressing, Figure 2 i) when comparing DSAD and EOAD. *PTN*, *COL25A1*, and *RASGRF1* expression, which were all elevated in protein abundance by mass spectrometry, were observed to be elevated in oligodendrocytes, astrocytes, and excitatory neurons, respectively, in DSAD compared with EOAD cases (Figure 2 m). We also detected reductions in *OTUD7B* and *RASGRF1* in oligodendrocytes (Figure 2 o), further underlining the diversity in gene expression across cell subtypes. When comparing DSAD with HA, we identified more changes in neuronal and astrocytic clusters, namely elevations in *NDE1, OTUD7B, PTN, SH38GRL3, STOM*, and *WASL* (Figure 2 p). Counter to what was identified using mass spectrometry, we also identified reductions in gene expression of *COL25A1* in a subtype of inhibitory neurons, *NLK* in oligodendrocytes, *NT5C* in microglia, and *PTN* in astrocytes. Proteins that we identified to be downregulated in our proteomics studies (Table 6) had both up and downregulation of gene expression in DSAD compared to EOAD and HA (Supplementary Figure 2). Overall, these data show a discordance between gene and protein expression.

### APOE abundance is increased at protein and transcript levels in DSAD compared to AD in the frontal cortex

Our proteomic and transcriptomic study indicated that trisomy 21 leads to raised APOE abundance in the brains of people with DSAD. APOE has a known key role in AD-related processes including amyloid-β clearance and aggregation, the immune response to pathology, tau pathology development, and the maintenance of cerebrovascular homeostasis [41]. The isoform variation of the protein is associated with an increased risk of developing AD in the general population [10, 16, 41]. Moreover, very limited data shows that this protein has a different abundance in DSAD. Thus, we focused on validating our observed APOE omics changes to provide new insight into DSAD.

Our initial data indicated that trisomy 21 may cause an increase in APOE protein in individuals with DSAD compared to individuals who are not trisomic for Hsa21. To validate this hypothesis, we quantified APOE abundance in two larger case series of human post-mortem samples, including both our discovery cohort (Table 1) and validation cohort A (Table 2), by western blot using two different anti-APOE antibodies which bind to epitopes in different parts of the APOE protein sequence (Polyclonal Goat, Calbiochem, 178479; Polyclonal Rabbit anti-APOE, C-terminus epitope, Sigma, SAB2701946). Consistent with our proteomic result, our western blot analysis using both antibodies showed that APOE abundance was significantly higher in DSAD compared to EOAD (Figure 3 a, b, e, f). To determine if a difference in the abundance of *APOE* transcript was observed between these cases in total tissue, we undertook RT-qPCR on total RNA isolated from the frontal cortex. Using this method, we found that *APOE* transcript was higher in DSAD and HA controls compared with EOAD (Figure 3 i).

**Figure 3.**
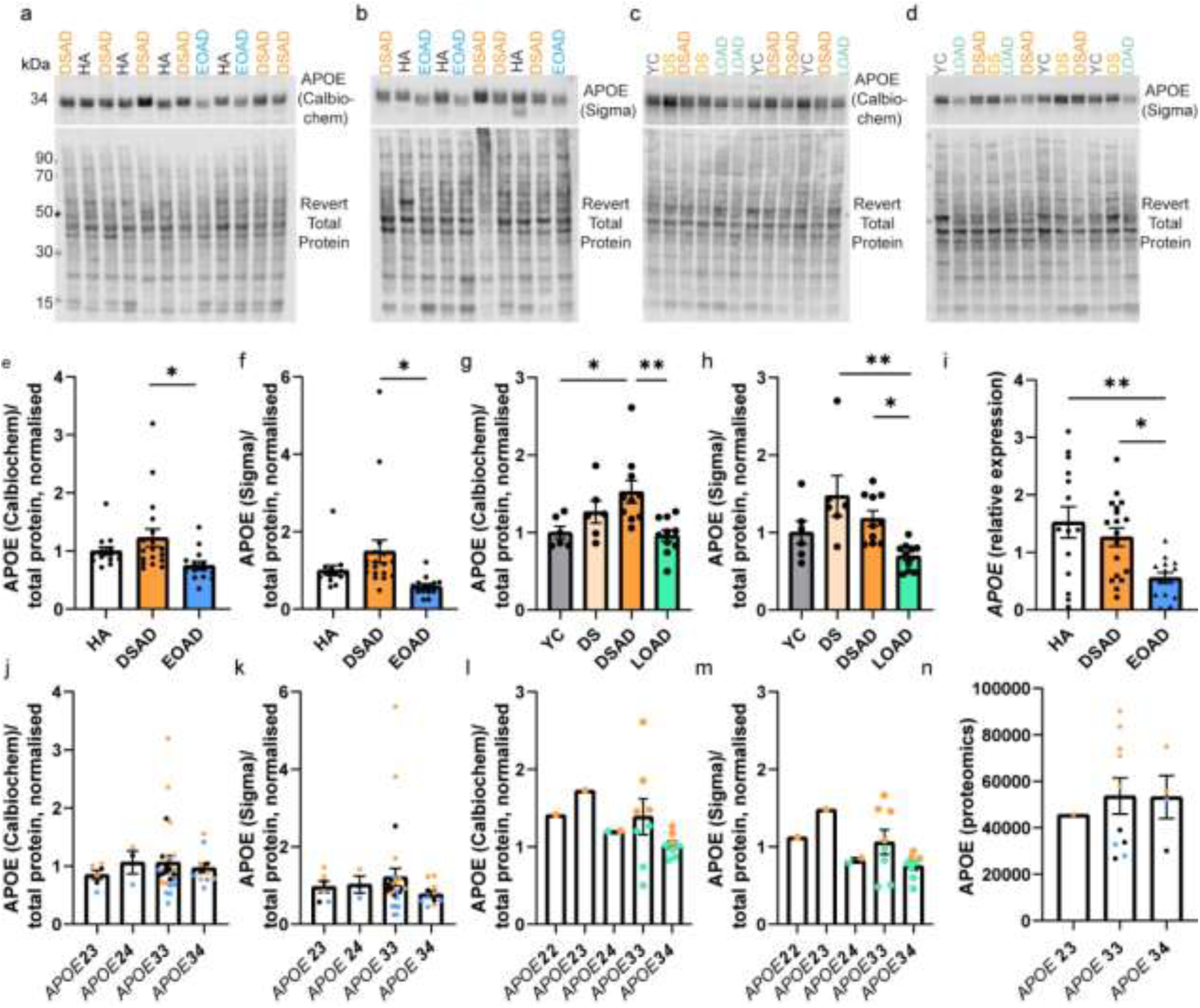
APOE abundance is increased in DSAD compared to matched cases of EOAD at protein and transcript level, independent of *APOE* genotype. (a-d) Representative western blots for APOE (Calbiochem, 178479), APOE C-terminal (Sigma, SAB2701946) and Revert 700 total protein stain (Licor bio, 926-11016) in frontal cortex samples from (a, b) discovery + validation cohort A (n=14 HA, n=18 DSAD, n=14 EOAD), and (c, d) validation cohort B (n=6 YC, n=6 DS, n=10 DSAD, n=10 LOAD). (e) Case type significantly alters APOE abundance (Calbiochem) (Univariate ANOVA F(2,43) = 4.381, p = 0.019), with APOE abundance significantly higher in DSAD than EOAD (post-hoc comparison with Bonferroni p = 0.015). (f) Case type significantly alters APOE abundance (Sigma) (Univariate ANOVA F(2,43) = 4.696, p = 0.014), with APOE abundance significantly higher in DSAD than EOAD (post-hoc comparison with Bonferroni p = 0.012). No effect of sex, age at death, or PMI was found for either antibody in the discovery and validation A cohorts. (g) Case type significantly alters APOE abundance (Calbiochem), in validation cohort B (Univariate ANOVA, F(3,28) = 5.541, p = 0.004, with APOE abundance significantly higher in DSAD than YC (post-hoc correction with Bonferroni, p = 0.033) and in DSAD than LOAD (p = 0.005). No effect of sex, age at death or PMI was found. (h) Case type significantly alters APOE abundance (Sigma), in validation cohort B (Univariate ANOVA, F(3,28) = 6.099, p = 0.003), with APOE being significantly higher in DS than LOAD (post-hoc comparison with Bonferroni, p = 0.002), and in DSAD than LOAD (p = 0.040). A significant interaction of age at death and case type was identified (F(1,22) = 6.169, p = 0.021), but no effect of sex or PMI were identified. (i) In qPCR from bulk frontal cortex tissue homogenate, case type alters *APOE* expression (Univariate ANOVA F(2,38) = 5.373, p = 0.009), with higher *APOE* expression in HA and DSAD than EOAD (post-hoc comparison with Bonferroni p = 0.004 and p = 0.033 respectively). APOE abundance by *APOE* genotype as detected by western blot (j, l) APOE (Calbiochem) (k,m), and APOE (Sigma), and by (n) mass spectrometry. (l, m) *APOE* genotype only available for DSAD and LOAD groups so only these cases included in analysis. *APOE* genotype had no effect on APOE abundance by western blot (j) (Calbiochem) in discovery and validation cohort A (Univariate ANOVA, F(3,42) = 0.333, p = 0.802), (k) (Sigma) in discovery and validation cohort A (Univariate ANOVA, F(3,42) = 0.624, p = 0.604), (l) (Calbiochem) in validation cohort B (Univariate ANOVA, F(5,26) = 1.142, p = 0.364), (m) (Sigma) in validation cohort B (Univariate ANOVA, F(5,26) = 1.488, p = 0.228), or by mass-spectrometry (Univariate ANOVA F(2,12) = 0.055, p = 0.947). Data expressed as mean ± SEM, *p<0.05, **p<0.01.

We also wanted to understand if increased APOE was caused by trisomy 21 alone, or due to an interaction of trisomy and AD. We therefore expanded our study to examine if APOE abundance was altered in the brain of young individuals with DS, prior to AD onset. Moreover, we wanted to determine if increased APOE in DSAD compared to EOAD was specific to the comparison to age-matched AD cases from the general population, or was also true for other types of AD, such as LOAD, which have a significantly later age of disease onset. Thus, we expanded our study to include cases in validation cohort B (Table 3). This included young DS cases (DS) (n = 6) and age-matched controls (YC) (n = 6), and cases of DSAD (n = 10) and LOAD (n = 10) which were not matched for age. We demonstrate that APOE abundance is significantly higher in DSAD than LOAD (Figure 3 c, d, g, h), supporting our initial findings that APOE is increased in DSAD compared to AD without DS. Moreover, we show that APOE abundance is significantly higher in DSAD than in YC (Calbiochem antibody only) (Figure 3 g).

Whether APOE is increased in the brains of young individuals with DS, before the onset of AD, is still unclear. No significant changes between DS and YC were detected using either antibody (Figure 3 g, h). However, we show that APOE abundance is higher in DS than in LOAD (Sigma antibody only), with values comparable to the DSAD group (Figure 3 h). Overall, we are underpowered for our frontal cortex YC and DS cases in this study, due to the low availability of these samples. To address this limitation, we measured APOE abundance in posterior cingulate cortex samples of YC and DS cases (n = 11 per group) (Supplementary Table 1). No significant difference in APOE abundance was identified between YC and DS cases using either antibody (Supplementary Figure 3 a-d). Whether an increase in APOE abundance is specific to DSAD cases or is only a region-specific phenotype in the frontal cortex requires further investigation.

We also had the opportunity to measure APOE abundance in the frontal cortex from an individual who had partial trisomy without the *APP* gene region [11]. As only one case of partial trisomy 21 was included, conclusions on APOE abundance in this individual cannot be made, however, APOE abundance in this sample was comparable to DS and DSAD (Supplementary Figure 4).

### Differences in APOE abundance between DSAD and EOAD are not the result of technical outliers

For western blots carried out on the discovery and validation A cohorts of HA, DSAD and EOAD samples, three cases had significantly lower total protein abundance than other cases. This resulted in three technical outliers (as measured by the ROUT method Q = 1%) for these western blots, two in the DSAD group, and one in the HA group, with higher abundances than other samples. To understand the effect of these technical outliers on our overall results, we repeated the statistical analysis removing the three cases. When removed from univariate ANOVA analysis, we continued to see a significant effect of case type on APOE abundance (Calbiochem) (Univariate ANOVA, F(2,40) = 5.005, p = 0.011) with DSAD being significantly higher than EOAD (post-hoc correction with Bonferroni, p = 0.010). A significant interaction of case type and sex was found (F(2,21) = 4.992, p = 0.017), but no effect of age at death, *APOE* genotype or PMI was found. We also continue to see a significant effect of case type on APOE abundance (Sigma) (Univariate ANOVA, F(2, 43) = 15.386, p < 0.001) with APOE being significantly higher in DSAD than EOAD (post-hoc correction with Bonferroni, p < 0.001), and APOE being significantly lower in EOAD than HA (post-hoc correction with Bonferroni, p = 0.011). A significant effect of PMI was identified (F(1,21) = 4.708, p = 0.042). No effect of sex, age at death or *APOE* genotype were identified. Thus, differences in protein abundance between cases did not alter our finding that APOE abundance is elevated in DSAD compared to matched cases of EOAD.

### APOE abundance is independent of APOE genotype

*APOE* genotype is known to influence AD risk, and our study contained a range of *APOE* genotypes. Thus, we examined the effect of *APOE* genotype on APOE abundance. No effect of *APOE* genotype was identified by western blot using either APOE antibody across cohorts, or by mass spectrometry proteomics (Figure 3 j-n). However, our proteomic study was underpowered to make pairwise comparisons between all *APOE* genotypes (Figure 3 n), as we only had one *APOE ε*2*ε*3 case, ten *APOE ε*3*ε*3 cases and four *APOE ε*3*ε*4 cases, thus further investigation is necessary to interpret the effect of *APOE* genotype on APOE abundance as detected by mass spectrometry. In our western blot study of validation cohort B, we did not have *APOE* genotype data on YC or DS cases, and so only DSAD and LOAD cases were included in the analysis. In these groups, only one *APOE ε*2*ε*2, one *APOE ε*2*ε*3 and two *APOE ε*2*ε*4 cases were included, and so we were underpowered to assess the effects of these genotypes on APOE abundance (Figure 3 l, m). However, the data we have indicates that the observed increase in APOE abundance in DSAD may not relate to *APOE* genotype.

### APOE abundance correlates with the abundance of APP, APP-CTFs, and amyloid-β, in the frontal cortex of DSAD and EOAD cases

We hypothesised that the increased abundance of APOE in the frontal cortex of individuals with DSAD was due to having an additional copy of a gene or genes on Hsa21. To identify which Hsa21-encoded proteins might contribute to this, we undertook correlation analysis of APOE and Hsa21-encoded proteins within our proteomic dataset. When all cases were included, APOE abundance significantly correlates with the abundances of APP, PDXK, S100B, CSTB and SYNJ1 (Table 7). To determine if the DSAD cases were driving the correlations we observed between these Hsa21 candidates and APOE abundance, we analysed only EOAD and HA cases (euploid cases), excluding the DSAD cases. In this sub-analysis, only the correlation between APOE and APP (Slope = 2.984, Pearson’s R = 0.8442, F(1,5) = 12.41, p = 0.0169), and APOE and S100B (Slope = 8.939, Pearson’s R = 0.7859, F(1,5) = 8.077, p= 0.0362) replicated (Table 7). Thus, we took these two proteins forward as lead Hsa21 candidates to investigate their role in increased APOE in DSAD compared with EOAD.

**Table 7:**
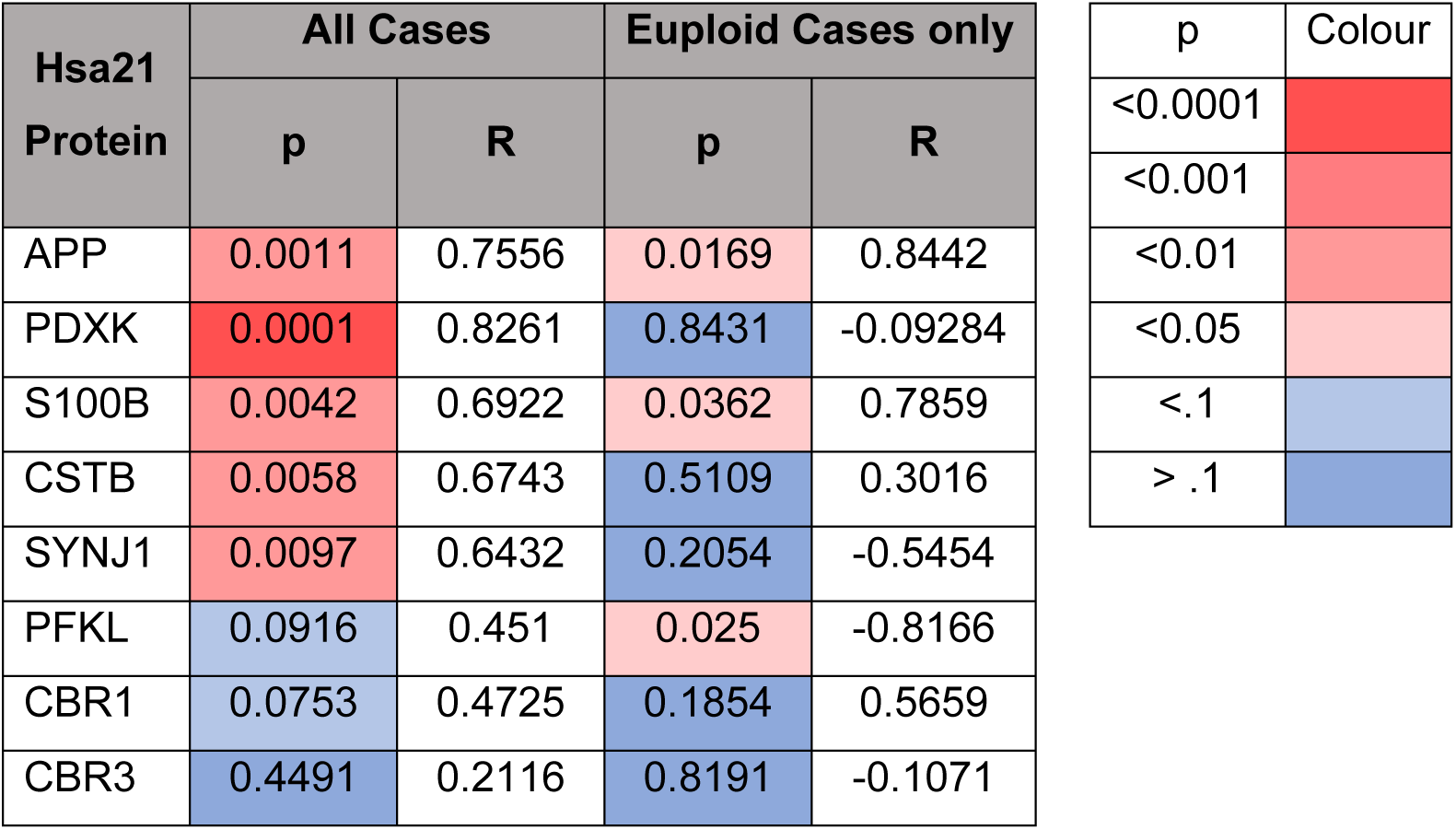
Correlation of the abundance of chromosome 21 encoded proteins and APOE in cases of HA, DSAD and EOAD frontal cortex. Correlation analysis (Pearson’s R and p-value) within the proteomic dataset between the abundance of APOE and all significantly altered Hsa21 encoded proteins in all cases (left) and only euploid (HA and EOAD) cases (right). APOE significantly positively correlates with APP and S100B, in both correlations. Red colour indicates a significant change, blue no significant change; intensity of colour indicates a lower p value.

We hypothesised that raised APOE in individuals with DSAD is the result of raised abundance of APP or/and S100B. We tested this by western blot and correlation analysis of APP and S100B with APOE abundance in our discovery cohort and validation cohort A samples. Consistent with our proteomic analysis and a previous report [55], we found APP abundance was increased in the DSAD frontal cortex (Figure 4 d, e). In contrast, we did not find a significant increase in S100B abundance in the frontal cortex of DSAD cases (Figure 4 a, b). Consistent with our proteomic data correlation analysis (Table 7), we found a significant positive correlation between APOE abundance and full-length APP. We also found that *APP* had increased expression in the same cell types as *APOE* in our snRNAseq transcriptomic study (Figure 3 m, n). However, in contrast to our proteomic data, by western blot we found a significant negative correlation between APOE and S100B abundance (Figure 4 c, h).

**Figure 4.**
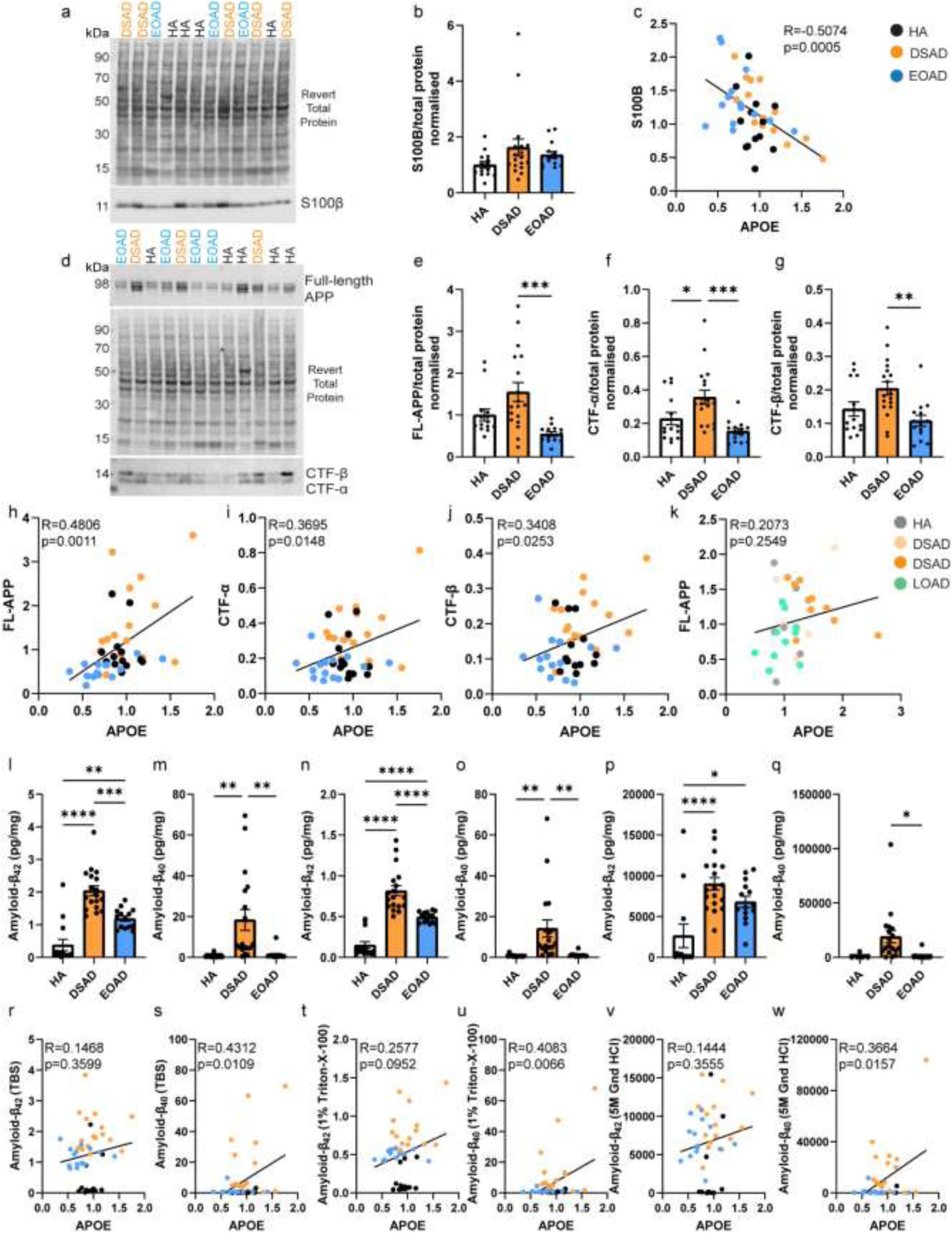
The abundance of APOE correlates with full-length APP, APP C-terminal fragments, and amyloid-β_40_ in frontal cortex. (a) Representative western blot for S100B (Abcam, ab41548) and Revert 700 total protein stain (Licor bio, 926-11016) in frontal cortex samples from the discovery cohort and validation cohort A. (b) Relative S100B abundance does not significantly differ between case types (Univariate ANOVA, F(2,38) = 2.188, p = 0.126). (c) S100B abundance negatively correlates with APOE abundance (Slope = -0.8321, Pearson’s R = -0.5074, F(1,41) = 14.21, p = 0.0005). (d) Representative western blot for APP (Abcam, Y188) showing bands representing full-length APP (FL-APP), APP-C-terminal fragment-α (CTF-α), APP-C-terminal fragment-β (CTF-β), and Revert 700 total protein stain (Licor bio, 926-11016) in frontal cortex samples from the discovery cohort and validation cohort A. (e) FL-APP abundance is significantly different between case types (Univariate ANOVA, F(2,38) = 7.155, p = 0.002), being elevated in DSAD compared to EOAD (post-hoc correction with Bonferroni, p < 0.0001), and tending to be elevated in DSAD compared to HA (p = 0.081). (f) CTF-α abundance differed between case types (Univariate ANOVA, F(2,38) = 7.025, p = 0.003) with elevated abundance in DSAD compared with EOAD (post-hoc correction with Bonferroni, p = 0.001) and DSAD compared with HA (p = 0.038). (g) CTF-β abundance differed between case types (Univariate ANOVA, F(2,38) = 5.493, p = 0.008) with elevated expression in DSAD compared with EOAD (post-hoc correction with Bonferroni, p = 0.003) and tending to be elevated in DSAD compared to HA (p = 0.087). (h) Correlation of FL-APP with APOE (Calbiochem) western blot abundance showed a significant positive correlation (Slope = 1.358, Pearson’s R = 0.4806, F(1,41) = 12.32, p = 0.0011). (i) Correlation of CTF-α with APOE (Calbiochem) western blot abundance showed a significant positive correlation (Slope = 0.2049, Pearson’s R = 0.3695, F(1,41) = 6.482, p = 0.0148). (j) Correlation of CTF-β with APOE (Calbiochem) western blot abundance showed a significant positive correlation (Slope = 0.1040, Pearson’s R = 0.3408, F(1,41) = 5.388, p = 0.0253). (k) Correlation of FL-APP with APOE (Calbiochem) abundance in validation cohort B did not show a significant relationship (Slope = 0.2387, Pearson’s R = 0.2073, F(1,30) = 1.347, p = 0.2549). MSD amyloid-β multiplex assay was used to quantify the abundance of (l-q) amyloid-β_42_ and amyloid-β_40_ in (l, m) soluble (Tris buffered saline), (n, o) membrane-associated (1% Triton-X100) and (p, q) insoluble aggregated (5M guanidine hydrochloride) fractions of frontal cortex from cases of HA, DSAD and EOAD (discovery cohort and validation cohort A). (l) Soluble amyloid-β_42_ abundance differed between case types (Univariate ANOVA, F(2,38) = 41.312, p < 0.0001), with significantly elevated levels in DSAD (post-hoc correction with Bonferroni p < 0.001) and EOAD (p < 0.001) than HA, and higher levels in DSAD than EOAD (p < 0.001). (n) Membrane-associated amyloid-β_42_ abundance differed between case types (Univariate ANOVA, F(2,38) = 45.256, p < 0.0001), with significantly elevated levels in DSAD (post-hoc correction with Bonferroni, p < 0.001) and EOAD (p < 0.001) than HA, and higher levels in DSAD than EOAD (p < 0.001). (p) Insoluble aggregated amyloid-β_42_ abundance differed between case types (Univariate ANOVA, F(2,38) = 11.322, p < 0.0001), with significantly elevated levels in DSAD (post-hoc correction with Bonferroni, p < 0.001) and EOAD (p = 0.016) than HA, and no difference between DSAD and EOAD (p = 0.281). (m) Soluble amyloid-β_40_ abundance differed between case types (Univariate ANOVA, F(2,38) = 7.215, p = 0.002), with significantly elevated levels in DSAD than HA (post-hoc correction with Bonferroni, p = 0.003), and EOAD (p = 0.004). (o) Membrane-associated amyloid-β_40_ abundance differed between case types (Univariate ANOVA, F(2,38) = 5.960, p = 0.006), with significantly elevated levels in DSAD than HA (post-hoc correction with Bonferroni, p = 0.006), and EOAD (p = 0.007). (q) Insoluble aggregated amyloid-β_40_ abundance differed between case types (Univariate ANOVA, F(2,29) = 3.339, p = 0.050), with significantly elevated levels in DSAD than EOAD (post-hoc correction with Bonferroni, p = 0.021). (r, t, v) No significant correlation was found between APOE (Calbiochem) abundance by western blot and amyloid-β_42_ in soluble or insoluble frontal cortex protein fractions. A significant positive correlation was identified between APOE (Calbiochem) abundance by western blot and amyloid-β_40_ in the (s) soluble (Slope = 20.83, Pearson’s R = 0.3664, F(1,41) = 6.359, p = 0.0157), (u) membrane-associated (Slope = 19.09, Pearson’s R = 0.4083, F(1,41) = 8.205, p = 0.0066) and (w) insoluble aggregated (Slope = 27840, Pearson’s R = 0.4312, F(1,32) = 7.307, p = 0.0109) frontal cortex fractions. Discovery and validation cohort A; n=14 HA, n=18 DSAD, n=14 EOAD. Validation cohort B; n = 6 YC, n = 6 DS, n = 10 DSAD, n = 10 LOAD. Data expressed as mean ± SEM, *p<0.05, **p<0.01, ***p<0.001, ****p<0.0001.

Full-length APP is processed into several fragments within the brain, including C-terminal fragments α (APP-CTF-α) and β (APP-CTF-β), amyloid-β_40_ and amyloid-β_42_ which accumulate in plaques in AD and DSAD. We, therefore, determined whether there was a relationship between these APP cleavage products and APOE abundance in our case series. For this analysis, we excluded the three technical outliers, for which samples had low total protein abundance. We first quantified the abundance of APP-CTF-α and APP-CTF-β by western blot and observed that both fragments had increased abundance in the frontal cortex of individuals who had DSAD, compared to both matched cases of EOAD (Figures 4 d, f, g). APP-CTF-α was also found to be elevated in DSAD compared with HA (Figures 4 f). This is likely the result of the additional copy of *APP* in individuals with DS and is consistent with a previous report [55]. We found that the abundance of full-length APP, APP-CTF-α, and APP-CTF-β, positively correlated with the abundance of APOE in the frontal cortex in cases of DSAD and EOAD (Figure 4 h-j). We also found these positive correlations with APOE (Sigma) by western blot (Supplementary Table 2). In validation cohort B, consisting of cases of DSAD and LOAD, we did not replicate the correlation between full-length APP and APOE abundance in the frontal cortex (Figure 4 k, Supplementary Figure 5). However, a positive correlation between full-length APP and APOE abundance in the young posterior cingulate cortex samples of YC and DS cases was observed (Supplementary Figure 3g).

Previous studies have shown that APOE is sequestered in amyloid-β plaques within the brain, including in individuals with DSAD [12, 3], thus we determined if the abundance of APOE correlated with the abundance of aggregated amyloid-β in our case series. To quantify the abundance of aggregated amyloid-β in our samples, we biochemically fractionated total protein into soluble, membrane-associated and plaque-associated/aggregated protein fractions. Consistent with the diagnosis of AD in the EOAD and DSAD cases, we found that the abundance of amyloid-β_42_ was raised in all three fractions in these case types compared with HA controls (Figure 4 l, n, p). Moreover, we found that in the soluble and membrane-associated fractions, the abundance of amyloid-β_42_ was higher in DSAD than in EOAD cases (Figure 4 l, n). We also found that the abundance of amyloid-β_40_ was increased in soluble and membrane-associated fractions isolated from cases of DSAD compared with both HA controls and cases of EOAD (Figure 4 m, o), likely caused by the increased production of this fragment due to trisomy 21. We found the insoluble aggregated amyloid-β_40_ was elevated in DSAD compared to EOAD but did not differ significantly between DSAD and HA cases, due to few HA cases having above threshold values for insoluble amyloid-β, and thus not being included in our statistical analysis because they were below the limit of detection (Figure 4 q). We found that amyloid-β_40_, in the soluble, membrane-associated, and aggregated fractions, significantly positively correlates with APOE abundance (Figure 4 s, u, w). However, no correlation between amyloid-β_42_ and APOE abundance was found in any fraction (Figure 4 r, t, v). These data indicate that APOE abundance is higher in DSAD than in EOAD and LOAD. Moreover, we find evidence that the abundance of APP and its processing products correlate with APOE abundance in our DSAD and EOAD case series.

### Increased APOE correlates with total tau abundance, but not phosphorylated tau

We hypothesised that increased APOE may contribute to the recently reported accelerated development of tau pathology in individuals with DSAD compared to AD in the general population [71]. To investigate this, we correlated the abundance of APOE with total and phosphorylated AT8 tau abundance in our case series. We did not find any significant changes in total tau or AT8 abundances in DSAD and EOAD compared to HA controls (Figure 5 a, b, d, e), perhaps due to the limited sensitivity of western blotting for this purpose. We found a positive correlation between APOE abundance and total tau (Figure 5 c). No relationship between APOE and pathological AT8 abundance was found in our case series (Figure 5 c, f). but note that all our cases were end-stage AD, so a possible role of elevated APOE earlier in the AD development cannot be ruled out.

**Figure 5.**
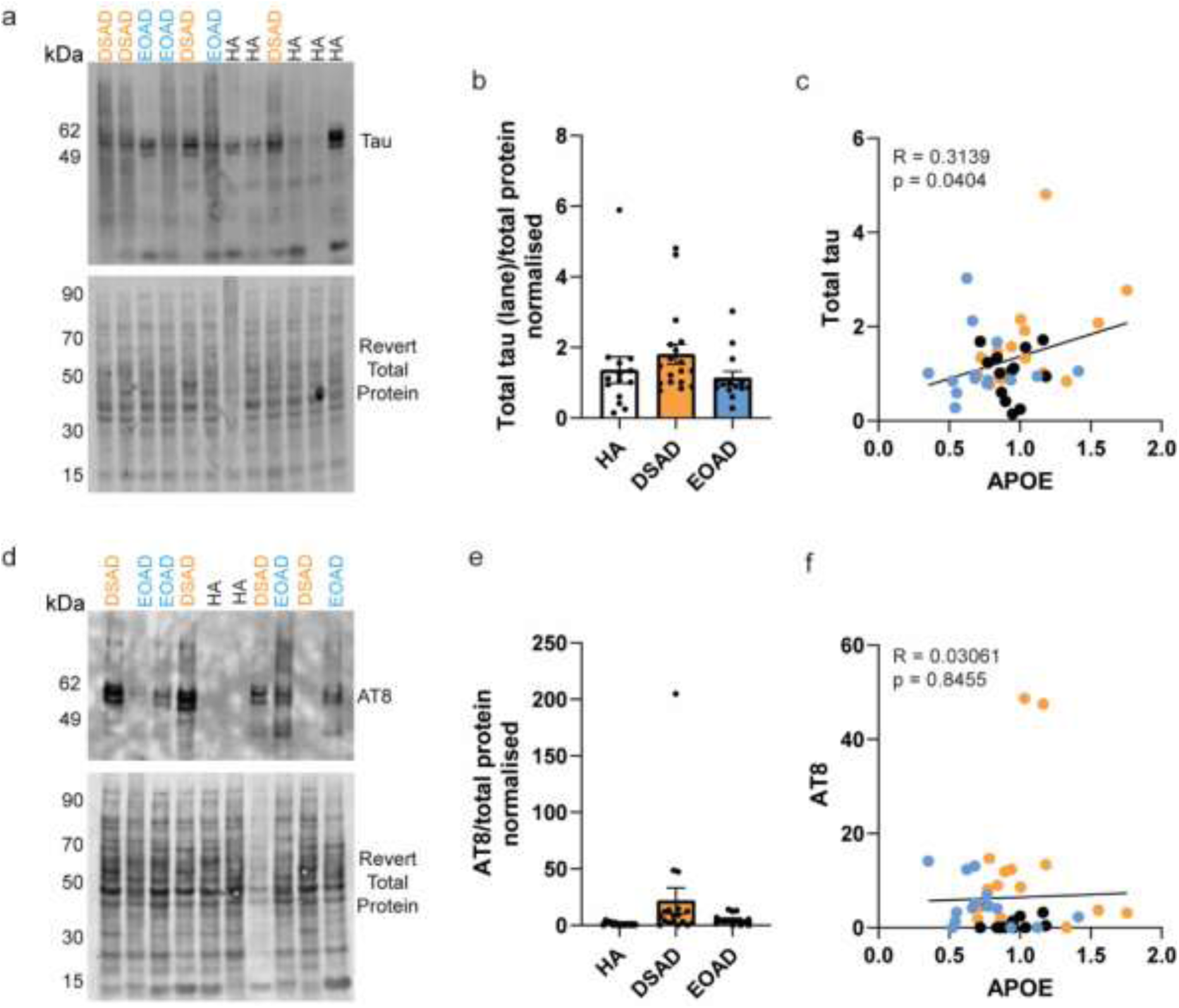
Total tau protein, but not AT8, correlates with APOE abundance. The abundance of (a, b) total tau (lane quantified), and (d, e) AT8 phosphorylated tau was quantified in frontal cortex by western blot and correlated (c, f) with the abundance of APOE to determine if there was a relationship between these proteins in the discovery cohort and validation cohort A. (b) No significant effect of case type was found on tau total abundance (Univariate ANOVA, F(2,38) = 0.867, p = 0.428). (c) A correlation between APOE and total tau abundance was observed (Slope = 0.9412, Pearson’s R = 0.3139, F(1,41) = 4.480, p = 0.0404). (d) No significant effect of case type was found on AT8 abundance (Univariate ANOVA, F(2,38) = 1.741, p = 0.189). (f) No correlation between APOE and AT8 abundance was observed (Slope = 1.165, Pearson’s R = 0.03061, F(1,41) = 0.03844, p = 0.8455). n=14 HA, n=18 DSAD, n=14 EOAD. Data expressed as mean ± SEM.

## Discussion

Here, we present evidence that trisomy 21 results in changes to the proteome and transcriptome in the frontal cortex of people with DSAD compared to demographically matched cases of AD from the general population. Identified changes between DSAD and EOAD represent the effect of trisomy 21 on the AD-associated proteome and transcriptome. These changes include not only the upregulation of transcripts and proteins encoded on Hsa21, such as *APP/*APP, but a wider dysregulation of the transcriptome and proteome beyond Hsa21, as exemplified by the increase in *APOE/*APOE abundance in DSAD. Moreover, we demonstrated that APOE is also raised in DSAD compared to LOAD in an independent cohort of DSAD frontal cortex samples, indicating that this is a generalizable result. Together, our data indicate that the proteome in DSAD differs from other forms of AD, and that understanding the underlying biological cause of these changes is important to develop effective and safe treatments for this common cause of early-onset dementia.

The cause of increased APOE in DSAD is not clear, however, our snRNAseq data indicate that this change may result from the action of an additional copy of a gene(s) on Hsa21 in astrocytes, pericytes and/or endothelial cells. Here, we also observed that the abundance of APOE correlates with the abundance of full-length APP, and its processing products APP-CTFs, and amyloid-β_40_ in cases of EOAD and DSAD. Consistent with previous reports [46, 48], we observe that *APP* transcription is elevated in many cell types in the brains of people with DSAD, including cell types in which we find *APOE* expression is also raised. A previous report has indicated that raised APOE leads to an increase in *APP* gene expression in preclinical models via an effect on the transcription factor AP1 [28]. Importantly here, we know that raised APP abundance in our DSAD cases is driven by the additional copy of *APP* on Hsa21, as indicated by both human-genetics studies and preclinical modelling [11, 51, 55]. Therefore, the potential relationship between these two proteins in our study likely reflects the effect of increased gene dosage of *APP* on APOE in DS rather than an effect of APOE on APP. We also found a correlation between all forms of amyloid-β_40_ (soluble, membrane-associated and plaque-associated) and APOE abundance in EOAD and DSAD. A previous report found no significant difference in the abundance of APOE in amyloid-β plaques in the hippocampus and surrounding entorhinal/temporal cortex, from cases of DSAD and EOAD [12]. These data indicate that the mechanism of increased APOE in DSAD is unlikely to result from the enhanced sequestration of the protein in amyloid-β plaques.

We also find a negative correlation between APOE and S100B, as detected by western blot, in DSAD and EOAD frontal cortex. This suggests that elevated levels of APOE in DSAD are unlikely to be the direct result of the additional copy of *S100B* encoded on Hsa21. S100B is principally expressed in a subset of astrocytes in the adult brain and is upregulated in AD and DSAD [21, 40, 54, 62], particularly in brain regions with high neuritic plaque load, such as the frontal cortex. S100B is a calcium binding signalling protein that has an important role in neurodevelopment and brain damage response [26, 45]. Further work is needed to understand the mechanistic cause of increased APOE in DSAD brain, including identifying the Hsa21 causal gene(s) and the cell type(s) responsible for the elevated levels of the proteins.

APOE is a lipid-transport protein that provides cholesterol and other lipids to neurons and glia to maintain cellular homeostasis in the brain [41]. In the context of AD, *APOE* has been shown to be upregulated in response to pathology in microglia undergoing a damage-associated immune response [37]. However, in contrast to this literature, we see the downregulation of APOE in microglia in DSAD compared to EOAD. Instead, our transcriptomic data indicates that the identified increase in APOE in DSAD compared to AD may result from the elevated expression of *APOE* in endothelial cells, pericytes, and subtypes of astrocytes in the frontal cortex of people who had DSAD compared with individuals from the general population who had EOAD. These cell types make up the neurovascular unit and the blood-brain barrier. Blood-brain barrier breakdown has been demonstrated to occur in AD in the general population [61] but whether the blood-brain barrier is disrupted in DS or DSAD is currently unknown. People with DSAD have a significant burden of cerebral amyloid angiopathy (CAA), with more severe CAA found than in sporadic AD [25, 38]. How CAA affects the integrity of the blood-brain barrier is still not known. It is thought that CAA severity may be related to the incidence of amyloid-related imaging abnormalities in individuals treated with anti-amyloid-β immunotherapies, but more research is needed to understand the mechanisms through which this may occur [23]. Whether *APOE* in this study is upregulated in cells of the neurovascular unit in response to vascular changes or injury, or via a cell endogenous mechanism due to trisomy 21, is unclear. Future research should investigate the role of APOE in neurovascular cell types in the context of DS.

In this study, we found no consistent evidence of a relationship between *APOE* genotype and APOE abundance in the frontal cortex. However, our study size is small and a larger study of further cases with greater representation of *ε*2 and *ε*4 *APOE* alleles is required to verify this finding. Notably, recent work has indicated that in DSAD, the effect of *APOEε*4 genotype on the age of onset of clinical features of disease and AD biomarkers is significant, but relatively modest compared with the effect of the genotype in LOAD, and may be limited to females [4, 20, 31]. Neuropathological evidence indicates that *APOEε4* results in higher amyloid-β deposition in people with DSAD [30]. However, recent PET imaging studies found no evidence that *APOEε4* influences amyloid-β load in individuals with DSAD [5].

We also observed an increase in the abundance of the cytokine pleiotrophin (PTN) in our proteomic study and an increase in *PTN* expression in excitatory neurons and oligodendrocytes. Elevated CSF PTN is associated with clinical AD [60] and our data indicate that this may result from increased expression in these cell types. Furthermore, changes to cytokine abundance in the brains of people with DS have been demonstrated, alongside perturbed neuro-immune biology [15, 43, 48, 65]. The study of the biology of this cytokine in DSAD may provide unique insight into its role in AD.

Recent natural history studies comparing the development of autosomal dominant AD (ADAD) and DSAD have highlighted evidence of accelerated AD progression in people who have DSAD. This includes a more rapid development of tau spread as measured by positron emission tomography and plasma phospho-tau217 [57, 66, 71], and enhanced rates of cortical thinning [36], after normalising for amyloid-β accumulation. These data indicate that differences in biology between DSAD and ADAD mediate disease progression. One study has highlighted that plasma GFAP (produced by astrocytes), raised in response to amyloid-β accumulation in the brain may contribute to the more rapid development of tau pathology [6]. Here, we undertook exploratory correlation analysis to investigate if raised APOE abundance DSAD was related to tau or phosphorylated tau abundance. Further research exploring the link between astrocytic APOE and disease progression in DSAD is warranted; given these data and recent preclinical work highlighting the importance of astrocytic APOE in disease development [52].

Overall, our observed changes to the proteome resultant from trisomy 21 may have implications for the development and treatment of AD in people with DS. The raised levels of APOE and PTN are consistent with a growing body of data indicating that the neuroimmune system differs in people with DS [15, 33, 43, 48, 50, 64, 65]. Our transcriptomic data indicate that differences in cell types of the neurovascular unit may contribute to altered biology in DSAD. As neuroimmune interactions are proposed to underlie the transition from pathology to disease in AD in the general population [9], understanding these differences is important to facilitate the safe and effective treatment of AD in individuals who have DS.

### Study limitations

Here we compare end-stage AD from individuals who had or did not have trisomy 21 to investigate how the additional copy of the Hsa21 may impact disease. In these tissues, substantial changes will have occurred, including changes to the abundance of cell types particularly as a result of neuronal loss. This may confound the bulk analysis of protein abundance presented here. Moreover, changes that occur during the earliest phases of disease will not be observed in our study, and thus the correlative relationships we observe here may not reflect all mechanisms important to disease development.

In this study we focused on matching between cases for demographic factors of Braak and Braak stage, age at death, sex and *APOE* genotype, thus we compared DSAD with cases of EOAD with unknown genetic cause. The underlying cause of disease development in these two case-types likely differs. DSAD is the result of the overproduction of APP and resultant increased abundance of amyloid-β_40_ and amyloid-β_42_. Whereas our data indicates that the EOAD cases studied here have unchanged APP and amyloid-β_40_ abundance, but elevated amyloid-β_42_ abundance compared to HA controls. AD in these cases is likely driven by a change in APP processing, rather than overproduction of APP. This different aetiology of disease may impact on the results of this study, independently of the effect of trisomy 21. To address this limitation, we validated our principal finding of raised abundance in DSAD compared to EOAD in an independent cohort of DSAD cases compared with LOAD, in which disease is proposed to be principally the result of impaired clearance of amyloid-β.

The proteomic method used here identified 2855 proteins, including 23 encoded by Hsa21, representing only around 10-15% of the human proteome, and thus because of this technical limitation, many differences in protein abundance are likely to not be observed within this dataset. Thus, in addition to the correlative relationship between APP and APOE we observe in our dataset, other genes on Hsa21 may also contribute to altered APOE abundance. Our snRNAseq transcriptomic study analysed the transcriptome of 89,649 nuclei from 16 cases, but nuclei recovery was less than 5000 for two of the four cases of EOAD in our dataset. In addition, we did not recover high numbers of nuclei for all cell types for all cases, which may bias the observed differentially expressed genes.

In this study, we used two independent APOE antibodies to validate our finding of increased APOE abundance in DSAD. The binding sequence of the Calbiochem antibody is proprietary, and the Sigma antibody binds within the C-terminus of the APOE protein. In our western blots on validation cohort B, we see varying results between these antibodies, with the Calbiochem antibody showing no difference between DS and LOAD samples, but the Sigma antibody showing that DS cases have a significantly higher abundance of APOE than LOAD. Validation cohort B has a relatively small sample size in the YC and DS groups, which may contribute to this difference and further data is required to determine if APOE abundance differs between case types.

Despite these limitations, we have robustly identified alterations to the important AD-associated protein APOE in the brains of individuals with DSAD compared to individuals from the general population with AD. Our datasets open new hypotheses whereby future work can understand the biological and clinical implications of altered APOE in DSAD. Moreover, these newly generated datasets will serve as a hypothesis-generating tool for other key questions in the field.

## Conclusion

In conclusion, using label-free mass spectrometry proteomics and snRNAseq transcriptomics, we have identified the dysregulation of Hsa21 and non-Hsa21 genes and proteins in the brain of people with DSAD compared to age-matched individuals with AD or healthy ageing. Our study design allowed us to identify unique trisomy 21-driven changes, including the upregulation of APOE in the brains of people with DSAD compared with euploid individuals with EOAD or LOAD. Our data indicate that increased APOE may be driven by cells of the neurovascular unit: astrocytes, endothelial cells and pericytes. Moreover, we observe a correlation between *APP* gene products and APOE in DSAD and EOAD. The consequences of elevated APOE in DSAD remain unclear, but this will be critical to understanding DSAD disease mechanisms. Our data provide new insight into DSAD biology and expose differences from AD in the general population. It will be important to address these differences when considering therapeutic interventions for AD-dementia in people with DS.

## Abbreviations

AD: Alzheimer’s disease
APOE: Apolipoprotein E
APP: amyloid precursor protein
APP-CTF: amyloid precursor protein C-terminal fragment
DS: Down syndrome
DSAD: Alzheimer’s disease in individuals with Down syndrome
EOAD: early-onset Alzheimer’s disease
HA: healthy ageing individuals
Hsa21: Human chromosome 21
PFA: paraformaldehyde
PMI: post-mortem interval
SEM: standard error of the mean

## Declarations

### Human tissue

The use of human tissues in this study was in accordance with the UK Human Tissue Act (2004). Samples for the discovery cohort and validation cohort A were supplied, anonymized by the Newcastle Brain and Tissue Resource (NBTR), Newcastle University, Newcastle, UK and the South West Dementia Brain Bank (SWDBB), Bristol University, UK, and had full research consent (REC 19/NE/0008) and (REC 18/SW/0029). This work was carried out under NBTR and SWDBB’s NHS REC Research Tissue Bank ethical approval. Validation cohort B, studied at UCI, were obtained through IRB approved protocols and consent.

### Consent to publish

Not applicable

### Supplementary Information

Supplementary Information 1 contains data relating to our mass spectrometry based proteomic study. Sheet 1 is a list of all proteins identified by mass spectrometry. Sheet 2 is a list of all significantly altered proteins between case types. Sheet 3 is a table of all Hsa21-encoded proteins identified by mass spectrometry. Sheet 4 is a list of all APOE peptides identified in this study. Supplementary Information 2 contains data related to our single-nuclei RNAseq study. Sheet 1 contains the cell cluster IDs and the number of nuclei recovered for each case. Sheets 2-4 contain lists of differentially expressed genes between DSAD and EOAD, DSAD and HA and EOAD and HA comparisons.

### Availability of data and material

The datasets generated during the current study are available from the corresponding author on reasonable request. The mass spectrometry proteomics data have been deposited to the ProteomeXchange Consortium via the PRIDE [49] partner repository with the dataset identifier PXD058779 and 10.6019/PXD058779. The data discussed in this publication have been deposited in NCBI’s Gene Expression Omnibus [13] and are accessible through GEO Series accession number GSE284141.

### Competing interests

The authors declare that they have no competing interests.

### Author Contributions

F.K.W applied for ethical permission for research. F.K.W., C.F., C.E.T., E.H. and K.M. conceptualised and designed the study. Material preparation and data collection was carried out by C.F., W.W.H., J.H., P.M., O.S.T., L.F.A., E.A., E. D., N.R., V.S. and C.E.T. Data analysis was carried out by C.F. and Y.B. The first draft of the manuscript was written by F.K.W and C.F., and all authors commented on versions of the manuscript. All authors read and approved the final manuscript.

### Funding Statement

F.K.W., is supported by the UK Dementia Research Institute (UKDRI-1014) through UK DRI Ltd, principally funded by the UK Medical Research Council. F.K.W is also supported by an Alzheimer’s Research UK Senior Research Fellowship (ARUK-SRF2018A-001 and ARUK-SRFEXT2022-001). C.F. was funded by an Alzheimer’s Society Ph.D. studentship awarded to F.K.W (AS-PhD-19a-007). This work was also supported by the Rosetrees Trust (MB2020\100003) awarded to F.K.W., an Alzheimer’s Research UK Network pump priming grant awarded to C.F., and a UCL Bouge Fellowship awarded to C.F.. The funders had no role in study design, data collection and analysis, decision to publish, or preparation of the manuscript. Tissue for this study was provided with support from the Brains for Dementia Research (BDR) programme, jointly funded by Alzheimer’s Society UK and Alzheimer’s Society. The SWDBB is further supported by BRACE (Bristol Research into Alzheimer’s and Care of the Elderly). E.H., E.D. and L.F.A. had funding support from NIH/NIA U19AG068054 and P30AG066519.

## Acknowledgments

We would like to thank the Newcastle Tissue and Brain Resource (NBTR), their donors and donor’s families for providing brain tissue for this study. We would like to thank the South West Dementia Brain Bank (SWDBB), their donors and donor’s families for donating brain tissue for this study. Tissue for this study was provided with support from the BDR programme, jointly funded by Alzheimer’s Research UK and Alzheimer’s Society, and BRACE. We would like to thank Carlo Sala Frigerio (University College London) for his contribution to the study design and technical assistance with the single-nuclei RNA sequencing experiment. We would like to thank Tara Spires-Jones (University of Edinburgh) and Selina Wray (University College London) for their invaluable technical advice for our validation studies. We would like to thank Amanda Heslegrave (University College London) for the use of equipment for this study. We acknowledge generous funding provided by The Charlotte and Yule Bogue Research Fellowships in Honour of Sir Charles Lovatt Evans and A.J. Clark, through the University College London Friends and Alumni Association, Inc. (UCLFAA). This work is supported by the NIHR GOSH BRC. The views expressed are those of the author(s) and not necessarily those of the NHS, the NIHR or the Department of Health. The authors are grateful to the participants for their gift of brain donation. For the purpose of Open Access, the authors have applied a CC BY public copyright licence to any Author Accepted Manuscript version arising from this submission.

## Notes

### Competing Interest Statement

The authors have declared no competing interest.

