## Supplementary data for "Apolipoprotein E abundance is elevated in the brains of individuals with Down syndrome-Alzheimer’s disease"

### Supplementary Information

Supplementary Table 1

| Case type and ID | Sex | Age at death (years) | Post-mortem interval (hours) |
|---|---|---|---|
| DS_1 | F | 1 | 28 |
| DS_10 | M | 25 | 24 |
| DS_11 | M | 25 | 22 |
| DS_12 | M | 19.87 | 14 |
| DS_13 | M | 15 | 14 |
| DS_3 | M | 2 | 17 |
| DS_4 | M | 23 | 15 |
| DS_5 | M | 24 | 24 |
| DS_7 | F | 39 | 12 |
| DS_8 | M | 19 | 26 |
| DS_9 | F | 3 | 11 |
| YC_10 | M | 25 | 21 |
| YC_11 | M | 19 | 14 |
| YC_12 | M | 25 | 23 |
| YC_13 | M | 23 | 18 |
| YC_14 | F | 47.3 | 5 |
| YC_15 | M | 2 | 25 |
| YC_2 | M | 1.72 | 25 |
| YC_3 | F | 2 | 24 |
| YC_7 | F | 39 | 17 |
| YC_8 | M | 19 | 24 |
| YC_9 | M | 24 | 24 |

**Supplementary Table 1 Case demographics for posterior cingulate cortex cases.** Cases were sourced from NIH NeuroBioBank (USA). No significant difference in age at death was identified between case types (Univariate ANOVA, $F(1,20) = 0.249$, $p = 0.623$). No significant difference in PMI was identified between case types (Univariate ANOVA, $F(1,20) = 0.204$, $p = 0.656$). YC = Young control, DS = Down syndrome, M = male, F = female.

Supplementary Table 2

| **Fraction and *APP* product** | | **p** | **R** |
|---|---|---|---|
| Total | APP-CTF-α | 0.0026 | 0.4473 |
| | APP-CTF-β | 0.0247 | 0.3422 |
| | FL-APP | 0.0001 | 0.6193 |
| 5M Gnd HCl | Amyloid-$\beta_{42}$ | 0.2878 | 0.1701 |
| | Amyloid-$\beta_{40}$ | 0.0041 | 0.4802 |
| 1% Triton | Amyloid-$\beta_{42}$ | 0.0047 | 0.4227 |
| | Amyloid-$\beta_{40}$ | 0.0015 | 0.4696 |
| TBS | Amyloid-$\beta_{42}$ | 0.0122 | 0.3791 |
| | Amyloid-$\beta_{40}$ | 0.0053 | 0.4178 |

| p | Colour |
|---|---|
| <0.0001 | |
| <0.001 | |
| <0.01 | |
| <0.05 | |
| <.1 | |
| > .1 | |

**Supplementary Table 2 APOE abundance (Sigma) by western blot correlates with APP and its processing products in discovery and validation cohort A.** As in Figure 4, correlation analysis was carried out between APOE abundance (Sigma, SAB2701946) by western blot and APP / APP-CTFs (Y188, Abcam) by western blot, and amyloid-β40 / amyloid-β42 by MSD assay. P-value and Pearson's R shown. Significant positive correlations are shown in red, with increased intensity of colour representing a more significant relationship.

### Supplementary Figures

Supplementary figure 1

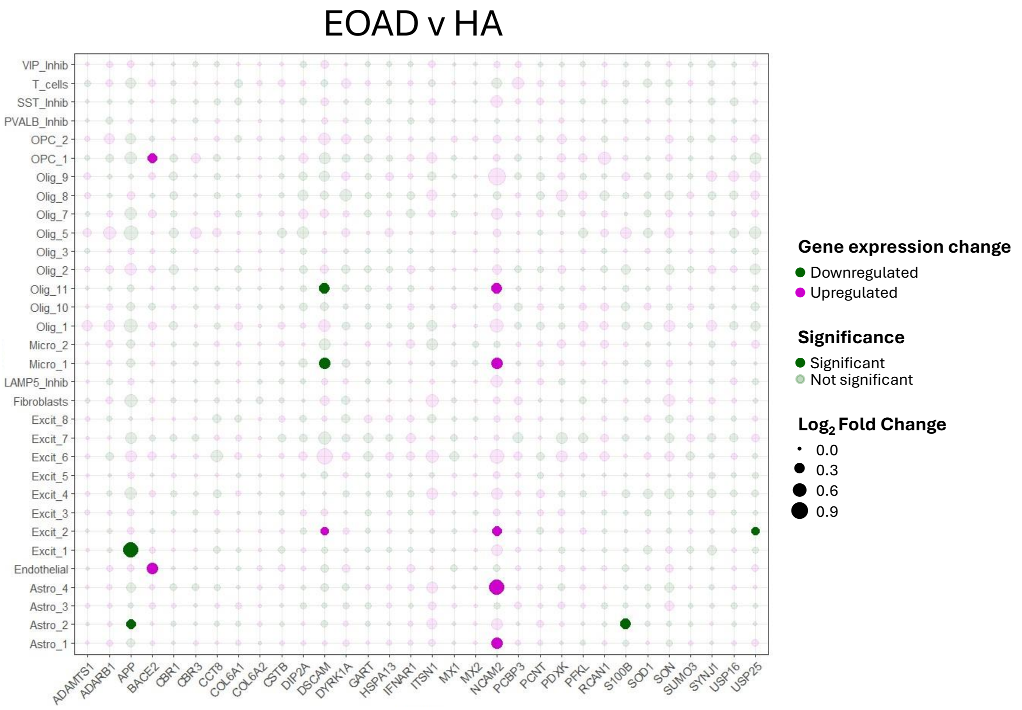


**Supplementary Figure 1 Differential gene expression analysis of Hsa21 genes in EOAD vs healthy ageing cohort (HA).** Hsa21-encoded genes identified in the proteomics dataset, and other commonly investigated Hsa21 genes, are dysregulated in few cell types in EOAD compared to HA.

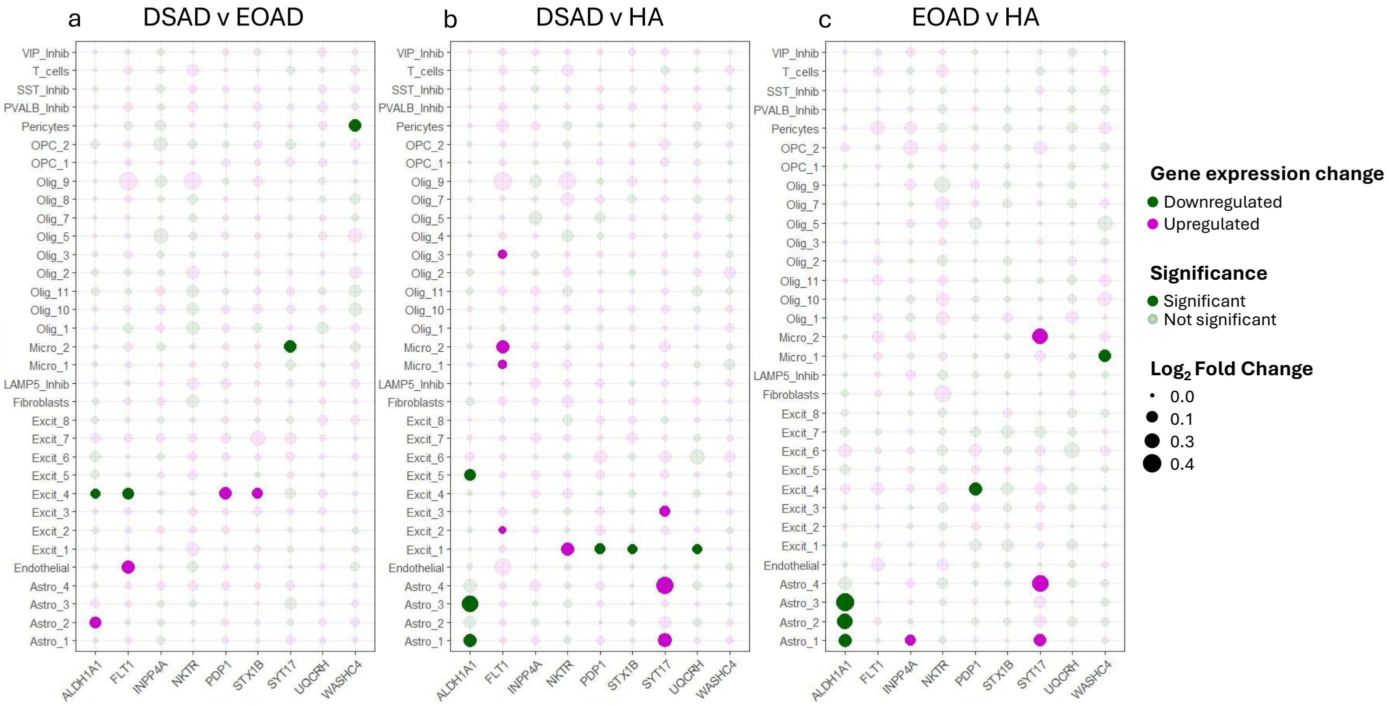


**Supplementary Figure 2 Single-nuclei RNA differential gene expression of top non-Hsa21 candidates that were downregulated in proteomics analysis.** (a-c) Non-Hsa21 genes which were significantly downregulated between case types in the proteomic study are represented in dot plots for DSAD againt EOAD (a), HA (b), and EOAD against HA (c).

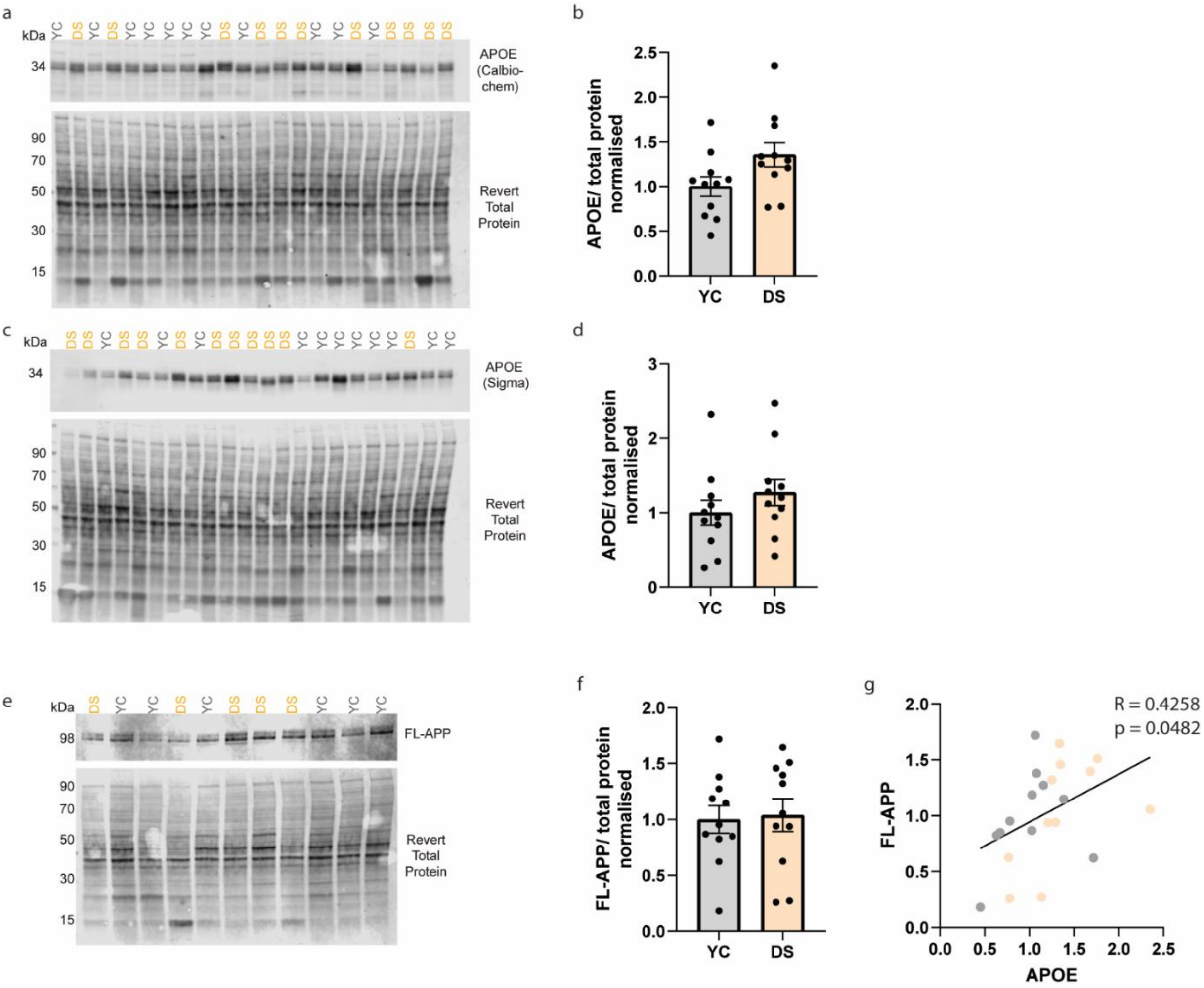


**Supplementary Figure 3 APOE is not different between young control and young DS posterior cingulate cortex samples.**

(a, c) Representative western blots for APOE (Calbiochem, 178479), APOE C-terminal (Sigma, SAB2701946) and Revert 700 total protein stain (Licor bio, 926-11016) in posterior cingulate cortex samples from young control (YC) and young DS (DS) (n=11 per group). (b) Case type has no significant effect on APOE abundance (Calbiochem) (Univariate ANOVA, $F(1,22) = 4.189$, $p = 0.054$). (d) Case type has no significant effect on APOE abundance (Sigma) (Univariate ANOVA, $F(1,22) = 1.231$, $p = 0.280$). (e) Representative western blot for APP (Abcam, Y188) and Revert 700 total protein stain (Licor Bio, 926-11016) in posterior cingulate cortex samples. (f) Case type has no effect on FL-APP abundance (Univariate ANOVA, $F(1,22) = 0.040$, $p = 0.844$). (g) APOE (Calbiochem) and FL-APP abundances positively correlate in young posterior cingulate cortex samples (Slope = 0.4278, Pearson's $R = 0.4258$, $F(1,20) = 4.430$, $p = 0.0482$). Data expressed as mean ± SEM.

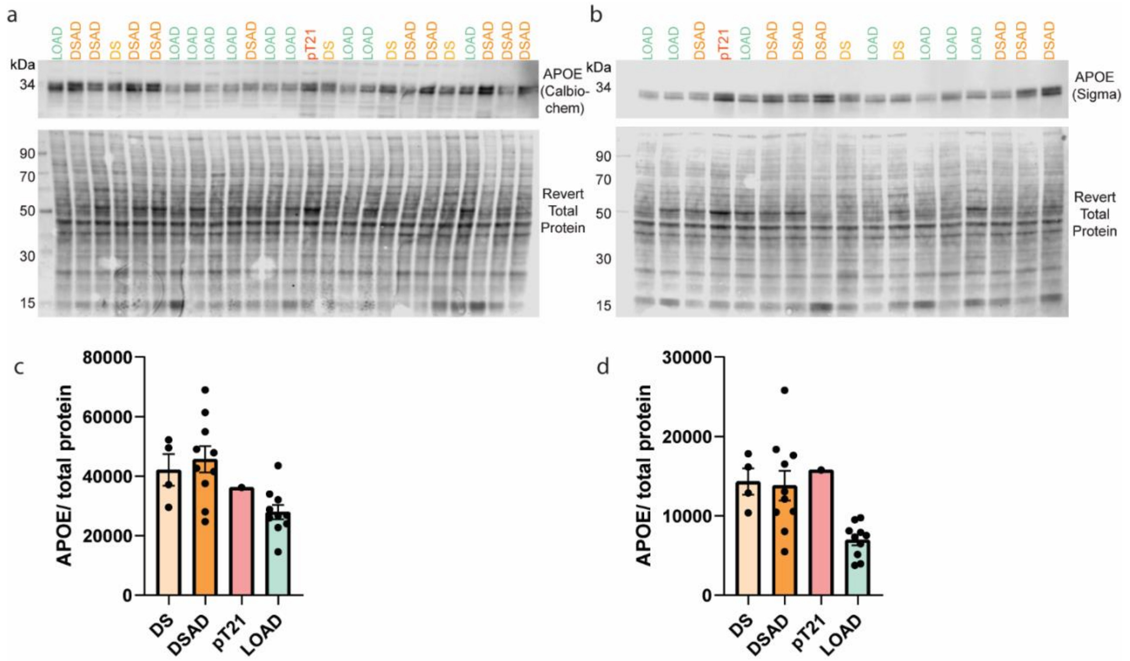


**Supplementary Figure 4 APOE abundance in frontal cortex of partial trisomy 21 case.** (a,b) Representative western blots for APOE (Calbiochem, 178479), APOE C-terminal (Sigma, SAB2701946) and Revert 700 total protein stain (Licor bio, 926-11016) in frontal cortex samples (validation cohort B) (n=4 DS, n=10 DSAD, n=1 pT21, n=10 LOAD). (c, d) As no young controls were included in this experiment, data is normalised to total protein only, and not internally normalised to young control values. Graphs for representative purposes only. Data expressed as mean ± SEM.
